# Ageing in a collective: The impact of ageing individuals on social network structure

**DOI:** 10.1101/2022.08.10.503309

**Authors:** Erin R. Siracusa, André S. Pereira, Josefine Bohr Brask, Josué E. Negron-Del Valle, Daniel Phillips, Cayo Biobank Research Unit, Michael L. Platt, James P. Higham, Noah Snyder-Mackler, Lauren J. N. Brent

## Abstract

Ageing affects many phenotypic traits, but its consequences for social behaviour have only recently become apparent. Social networks emerge from associations between individuals. The changes in sociality that occur as individuals get older are thus likely to impact network structure, yet this remains unstudied. Here we use empirical data from free-ranging rhesus macaques and an agent-based model to test how age-based changes in social behaviour feed up to influence: (1) an individual’s level of indirect connectedness in their network; and (2) overall patterns of network structure. Our empirical analyses revealed that female macaques became less indirectly connected as they aged for some, but not all network measures examined, suggesting that indirect connectivity is affected by ageing, and that ageing animals can remain well integrated in some social contexts. Surprisingly, we did not find evidence for a relationship between age distribution and the structure of female macaque networks. We used an agent-based model to gain further understanding of the link between age-based differences in sociality and global network structure, and under which circumstances global effects may be detectable. Overall, our results suggest a potentially important and underappreciated role of age in the structure and function of animal collectives, which warrants further investigation.

## Introduction

The costs and benefits of an animal’s behaviour toward members of their collectives can shape lifespans, life history evolution, and the pace of senescence. For example, the support that cooperative breeding species receive from non-breeding helpers may lead to longer lifespans than those of related solitary species [1]. The transfer of enhanced ecological knowledge to close relatives during collective foraging may have contributed to the evolution of reproductive senescence and prolonged post-reproductive lifespan (i.e., menopause) in female killer whales [2– 4]. The formation of strong relationships between group mates has also been linked to enhanced longevity in a range of group-living taxa [5]. It is therefore clear that living in collectives can alter the adult ageing process. However, an important question is whether ageing, in turn, influences the behaviour of adults in collectives, and ultimately the structure of collectives themselves.

Growing evidence suggests that older adults differ in their social behaviours and social relationships from young adults [6–11]. One pattern that seems to be emerging across taxa is that older adults interact with fewer individuals than do younger adults, concentrating social relationships on close associates and kin [6,12–15]. Given that social networks are an emergent feature of association rules between individuals [16], shifts in patterns of social behaviour with age might not only affect who ageing individuals associate with directly (i.e., their direct connectedness), but could also affect higher-order network structure. Age-based changes in social behaviour could scale up to alter an individual’s connections to the partners of their social partners (their indirect connectedness [17]) as well as the overall structure (“topology”) of the social network, both of which can have consequences for disease transmission [18–23], information transfer [24–26], the cohesive movement of groups [27,28], and many other eco-evolutionary dynamics [29,30]. Yet little attention has been given to understanding the impact of social ageing for the polyadic social world or the structure of the collective.

Understanding how ageing shapes an individual’s indirect connectedness may be particularly relevant as such connections are tightly linked to processes that can directly influence fitness, including those described above. Declines in indirect connectedness with age may help limit exposure to disease [20–22] which might be beneficial in aged animals experiencing immunosenescence, but simultaneously could inhibit the transfer of important socio-ecological information [26,31] which could exacerbate pre-existing fitness declines in old age. In some cases, indirect connectivity may be an even more important predictor of fitness than direct connections dyadic associations [17,32]. Therefore, changes in indirect connections may be a particularly important component of the social ageing phenotype to investigate. Recent work has offered some glimpses into how measures of indirect connectedness can differ between young and old adults. In marmots and Barbary macaques, older adults have measures of indirect connectedness that suggest they have partners who are themselves not well connected [9,33], (but this is not the case in rhesus macaques, see [34,35]). Older marmots and rhesus macaques are less effective at reaching disparate nodes in the network compared to younger adults [34–36]. In contrast, older adults are more strongly embedded in cliques or clusters in their networks compared to younger adults in marmots [9], but not in either Barbary or rhesus macaques [33,35]. However, most research to date has compared differences in measures of indirect connectedness among adults of different age classes (e.g., old versus young), but has lacked the longitudinal data required to quantify how the social positioning of individuals changes across their lifetimes (c.f., [9]). Such longitudinal analyses have the potential to reveal important patterns that might otherwise be masked by differences between individuals or cohorts [37]. Tracking within-individual changes in measures of indirect connectedness is essential for more firmly placing changes in sociality across the lifespan within the larger ageing phenotype and therefore understanding the causes and consequences of these patterns of social ageing [38].

Populations composed of a greater proportion of older (or younger) adults may also be structured in meaningfully different ways, and this could affect important processes such as communication and cooperation. For example, the loss of old individuals through age-related disease or trophy hunting can disrupt intergenerational flow of accumulated social and ecological knowledge, impeding collective movement and the ability to locate critical resources [4,39–42]. The age structure of a group can also regulate the behaviour of younger individuals [43,44], influencing aggression rates and social cohesion [45]. The impact of diminished cohorts of younger individuals on overall network structure is less well understood, but likely to have repercussions for network connectivity and cohesiveness given that younger adults are more socially active in many populations [8,10,12–14]. For example, the simulated removal of juvenile killer whales led to networks that were more fragmented than when random individuals were removed, suggesting an important role of young individuals in maintaining network cohesion [46]. Despite the established ecological and evolutionary importance of network structure [29,30] the underlying drivers of variation in network structure remain understudied [47–50]. Ageing, as an important process underlying patterns of individual-level variation in sociality, might therefore provide a window into how simple processes can generate complex network structures [51,52].

Using both empirical data and a theoretical model, we explore how social ageing of individuals relates to measures of indirect connectedness and overall network structure in a group-living primate, the rhesus macaque (*Macaca mulatta*), which is an emerging model in social ageing research [38,53]. As female rhesus macaques grow older, they show clear changes in their patterns of direct connectedness: they reduce the size of their social networks and focus their social effort on a few important partners including close kin [6]. Despite this, females do not reduce the rate at which they engage in social interactions as they age, indicating that although their networks get smaller, older females continue to invest the same amount of time into fewer relationships [6].

Given these previously established changes in direct connections, here, we set out to test if age-based changes in social behaviour relate to measures of an individual’s level of indirect connectedness [17,54], and if they scale up to influence network structure as a whole. Individual-level social network metrics, including measures of direct and indirect connectedness, relate to underlying, putatively simple, social differences or processes, such as individual-level variation in general sociability or reassociation tendency [51]. For example, an individual’s general tendency to be sociable can be intuitively quantified using a direct network metric (e.g., strength, or the sum of the weights of an individual’s ties to their partners), or using an indirect metric (e.g., weighted eigenvector centrality, which measures how well-connected an individual is to its partners and how well connected those partners are to others [51]). Drawing on this idea that underlying processes can predict an individual’s position in the network, and on the age-based changes in direct connections we have previously documented in rhesus macaque females [6], we made predictions for four common measures of indirect connectedness: eigenvector centrality, betweenness, closeness, and clustering coefficient (Table 1). Adult female rhesus macaques maintain their strength of grooming ties to others as they age, despite reducing their number of partners [6]. We therefore predicted that weighted eigenvector centrality (a measure of overall connectedness in the network) would also remain stable with age, as by retaining strong connections to some of their partners, females therefore (indirectly) retain connections to the partners of their partners. We also previously found that as female rhesus macaques age, they increase their likelihood of interacting with certain partners, especially their kin [6]. That is, ageing females increase both their tendency to reassociate with others, and their tendency to interact with someone from their own (kin-based) sub-group. Individuals with higher within-group association tendencies mix less widely in their networks and those with greater reassociation tendency are less likely to associate with new individuals and connect disparate parts of the network [51]. Therefore, we predicted that as females aged, they would have lower measures of betweenness (capacity for linking discrete clusters in a network) and closeness (capacity to reach others or be reached), but a higher clustering coefficient (cliquishness).

**Table 1.** Predictions for how indirect measures of connectedness are expected to change with age in female rhesus macaques from Cayo Santiago and how global network metrics are expected to change with increasing proportions of old individuals in the population.

| Network Metric | Definition | Prediction | Rationale |
| --- | --- | --- | --- |
| **Indirect network metrics** |  |  |  |
| **Eigenvector centrality** | Measures how well connected an individual is in the network to individuals who are themselves well connected. | Remain stable with increasing age | Females retain strength of connections to partners as they age. |
| **Betweenness** | Calculates the number of shortest paths between others in a network that pass through an individual. Individuals with low betweenness have low capacity for linking discrete clusters in the network. | Decrease with increasing age | Females increase their probability of interacting with strong, stable partners and kin. Therefore, females increase their tendency to reassociate with existing partners and have higher within-group association tendencies as they age. |
| **Closeness** | Measures the inverse distance to all other individuals in the network. The lower an individual's closeness score, the more difficult it is for them to reach others or be reached. | Decrease with increasing age |  |
| **Clustering** | Measures cliquishness or subgrouping. A high clustering coefficient indicates that an individual's partners are highly connected to each other. | Increase with increasing age |  |
| **Global network metrics** |  |  |  |
| **Mean degree** | The average number of ties that each individual has to others in the network. | Decrease with increasing proportion of old individuals | Older females have fewer social partners. |
| **Diameter** | Measures the overall connectedness of the network. Networks with a larger diameter are less connected. | Increase with increasing proportion of old individuals |  |
| **Transitivity** | The degree of clustering in the network. | Increase with increasing proportion of old individuals | Older females cluster more with kin. |

To determine if age-based changes at the individual level can result in changes in overall network structure we used empirical data from 19 networks of female rhesus macaques to test how variable proportions of old individuals in a network was related to three common measures of global network structure: mean degree, network diameter and transitivity (Table 1). We predicted that networks with a greater proportion of old individuals would be more sparsely connected due to older animals having fewer social partners (i.e., have lower mean degree and network diameter). Given greater kin clustering with age, we expected networks with more old individuals would be more clustered (i.e., have higher transitivity). Finally, to help inform our empirical findings and better understand the link between age distribution and global structure, we built an agent-based model (parameterized using information from our empirical data) to simulate how different proportions of old individuals would be expected to affect network structure under simplified conditions where everything else is equal. Our results provide a first step to understanding how and when individual changes in social tendencies with age might scale up to detectable effects on global network structure, offering important insight into the consequences of demography for the structure and function of collectives.

## Methods

### Study population and data

Data used in this study were collected on a well-studied population of rhesus macaques on the island of Cayo Santiago, off the southeastern coast of Puerto Rico. The current population is maintained by the Caribbean Primate Research Center (CPRC) and is descended from 409 macaques that were introduced to the island from India in 1938. The animals are food supplemented and provided with *ad libitum* access to water. There is no medical intervention, and so the major causes of death at this predator-free site are illness and injury [55,56]. The CPRC staff collect demographic data five days per week and thus track dates of birth and death of all individuals with a high degree of accuracy.

Rhesus macaques are highly social cercopithecine primates that live in matrilineal kin-groups and exhibit clearly differentiated social relationships with kin-biased affiliation [55,57,58]. At six years old, females are deemed adults [59] and previous research on the macaque population of Cayo Santiago has shown that, for females that survive to reproductive age, the median lifespan is 18 years with a maximum lifespan of about 30 years [53,60]. Female rhesus macaques have a strict dominance hierarchy with maternal rank inheritance and youngest ascendency [61]. Patterns of social interactions and social attention vary between young and old adults [6,34,62]. Female macaques show clear evidence of within-individual declines in the number of grooming partners with age, although the amount of time spent giving and receiving grooming remains constant across adulthood [38].

For this study, subjects were mature adult females 6 years and older [59] from 6 naturally formed mixed-sex social groups. We used data collected between 2010-2017, a time period for which we had detailed behavioural data from which to estimate social networks. We collected behavioural data between 07:30 and 14:00, which are the working hours of the field station, using 10-min focal animal samples and recording all behaviours continuously [63]. We balanced data collection to ensure equal sampling of individuals throughout the day and over the course of the year, resulting in approximately the same number of focal samples per individual per year.

For these analyses we used grooming interactions to build our networks, given the clear age-based changes in grooming associations previously demonstrated in this system [6]. Grooming behaviour was collected by recording the duration of a grooming bout along with the identities of the interactants and the direction of grooming. We focused only on interactions between adult females (≥ 6 years old) and did not include interactions with infants (<1 year old), juveniles (2-3 years old), or sub-adult females (4-5 years old). We also did not include interactions with males as we wanted to avoid capturing changes in socio-sexual behaviour with age. We established dominance ranks for all females in a given year using observed win-loss interactions (as per [64,65]). Rank was assigned as “high” (≥ 80% of other females dominated), “medium” (50%-79% of other females dominated) or “low” (≤ 49% of other females dominated).

### Social networks

We built 19 grooming networks including all adult females (≥ 6 years old) from the following group-years (group F 2010-2017; group HH 2014; group KK 2015; group R 2015-2016; group S 2011; group V 2015-2016), with data availability based on the focus of projects over time given limited person power. We used weighted network metrics, as these are more robust and provide higher resolution than binary measures [66]. In these weighted networks, edges represented the undirected rate of grooming between a pair of individuals (seconds of grooming/total number of hours that both individuals were observed in focal animal samples). We note that using undirected grooming rates may fail to capture some finer nuances of age-based variation in social behaviour if, for instance, older individuals were likely to give less grooming than they received [61]. However, our most recent within-individual analyses suggest that while the number of grooming partners does decline as females age, both the amount of grooming given and received remains constant [6]. Therefore, the use of undirected grooming rates to build networks should provide a comprehensive picture of how individual connectedness to the wider network changes with age, without missing essential changes in the form that the connectedness takes. The average size of our networks was 50.7 (± SE = 3.9) adult females. All network metrics were calculated in R (version 4.2.0; [67]) using the igraph package (version 1.3.1; [68]).

### Empirical analyses

All empirical models were fitted in a Bayesian framework with different error structures and random effects dependent on the data analysed (see below). We conducted all analyses using R (version 4.2.0; [67]) and fitted all models in the Bayesian software STAN [69] using the brms package (version 2.17.0; [70]). All fixed effects were given weakly informative priors (see Supplementary Information for more details). We ran all models for 10,000 iterations across two chains with a warm-up period of 2,000 iterations. We assessed model convergence by examining trace plots to assess sampling mixing and by ensuring Rhat = 1. We considered estimates of fixed effects to be significantly different from zero when the 95% credible intervals of the posterior distribution did not overlap zero.

### Investigating the relationship between age and indirect connectedness

For these analyses we set out to test how individuals’ level of indirect connectedness to their social network changed from prime adulthood into later life. In our dataset, the median age of adult females was 10-years and previous studies in this system have shown that individuals aged 10 and beyond show clear evidence of physical [71–74], immunological [75], reproductive [71], and social ageing [6]. To capture later-life changes in indirect connectedness we focused our analyses on individuals aged 10 and older, in line with previous ageing studies [6]. Females in this analysis therefore ranged between 10 and 28 years old, although, to be clear, all measures of indirect connectedness for these females were extracted from the networks including all adult females (≥ 6 years old; see above). We had 563 macaque years of data over 204 unique females, with an average of 2.8 years of data per individual (range: 1-8 years; Fig. S1). Because there was variation in the age-ranges over which individuals were sampled, we used a within-individual centering approach to capture changes in indirect connectedness across individuals’ lifespans [76]. Briefly, following the methodology of van de Pol & Wright (2009) we split our age term into a between-individual effect (calculated as the mean age of an individual across all observations; hereafter called average age) and a within-individual effect (calculated as the deviation of an individual’s age from their mean age; hereafter called within-individual age). This within-individual age term was our primary variable of interest and reflects how an individual’s deviation from its age affects its indirect connectedness in the network (see [6] for a more detailed description of these methods).

We fitted four models with our four response variables of interest: eigenvector centrality, betweenness, closeness, and clustering coefficient. All statistical models included average age and within-individual age as continuous fixed effects. Given the strict dominance hierarchy exhibited by female rhesus macaques [61], we assessed whether social status affected the change in an individual’s measures of indirect connectedness with age [6] by fitting an interaction between rank and within-individual age in all models. We removed the interaction when not significant. We included individual ID, group, and year, as random effects to account for repeated observations and to capture any variation in indirect connectedness measures that might be due to differences between individuals, groups, or years. We also fitted within-individual age as a random slope over individual ID to capture any among-individual variation in the change in indirect connectedness with age. We had no biological reason to expect non-linearities in the relationship between age and measures of indirect connectedness given that we were looking at changes from prime adulthood to old age and we’ve previously found a linear relationship between direct measures of connectedness and age [6]. Nevertheless, we fitted a model with quadratic terms for within-individual age and average age and compared that model to the model with only linear age terms using leave-one-out cross-validation in the brms package (version 2.17.0) [70]. The quadratic terms never improved the model fit and so were not considered further. For the eigenvector centrality model, we logit-transformed the response variable to improve model fit and fitted a model with a Gaussian error distribution. For betweenness we fitted a model with a zero-inflated Poisson distribution. For closeness and clustering coefficient we fitted models with a Beta error distribution and a zero-one-inflated Beta error distribution, respectively.

### Investigating the relationship between age distribution and network structure

Our analyses quantifying changes in indirect connectedness with age focused on females aged 10 and older to capture individual changes in social network position from prime age and beyond. Here we were interested in how the age distribution of a group was linked to network structure. For these analyses we therefore used our complete networks including all adult females aged 6 and older. We had three response variables of interest (three measures of global network structure) and fitted a separate model for each: mean degree, diameter, and transitivity. To ask if age distribution relates to network structure, we included the proportion of old individuals as a continuous predictor in all our models. Since 18 is the median age of death in this population and maximum lifespan is about 30 years, we considered individuals above 18 to be “old” [52,61].

The proportion of old females in our 19 networks ranged from 0.03 to 0.19 (Fig. 1; Fig. S2). Given that rhesus macaques live in matrilineal groups with kin-biased behaviour [53,77], we included the average relatedness of the network as a continuous covariate to account for differences between groups in general relatedness. We also included network density (calculated as the number of existing ties in the network divided by the number of possible ties) as a continuous fixed effect in the models of diameter and transitivity to account for the fact that density can drive variation in other global network metrics. We included year as a random effect to account for yearly variation that might affect network structure. For the diameter model, we log-transformed the response variable to improve model fit. We fitted all models with a Gaussian error distribution. To account for a potential non-linear relationship between age distribution and network structure we fitted all models with a quadratic term for proportion of old individuals and compared that to the model with only linear terms using leave-one-out cross-validation in the brms package (version 2.17.0; [70]). The quadratic terms never improved model fit and so were not considered further.

**Figure 1.**
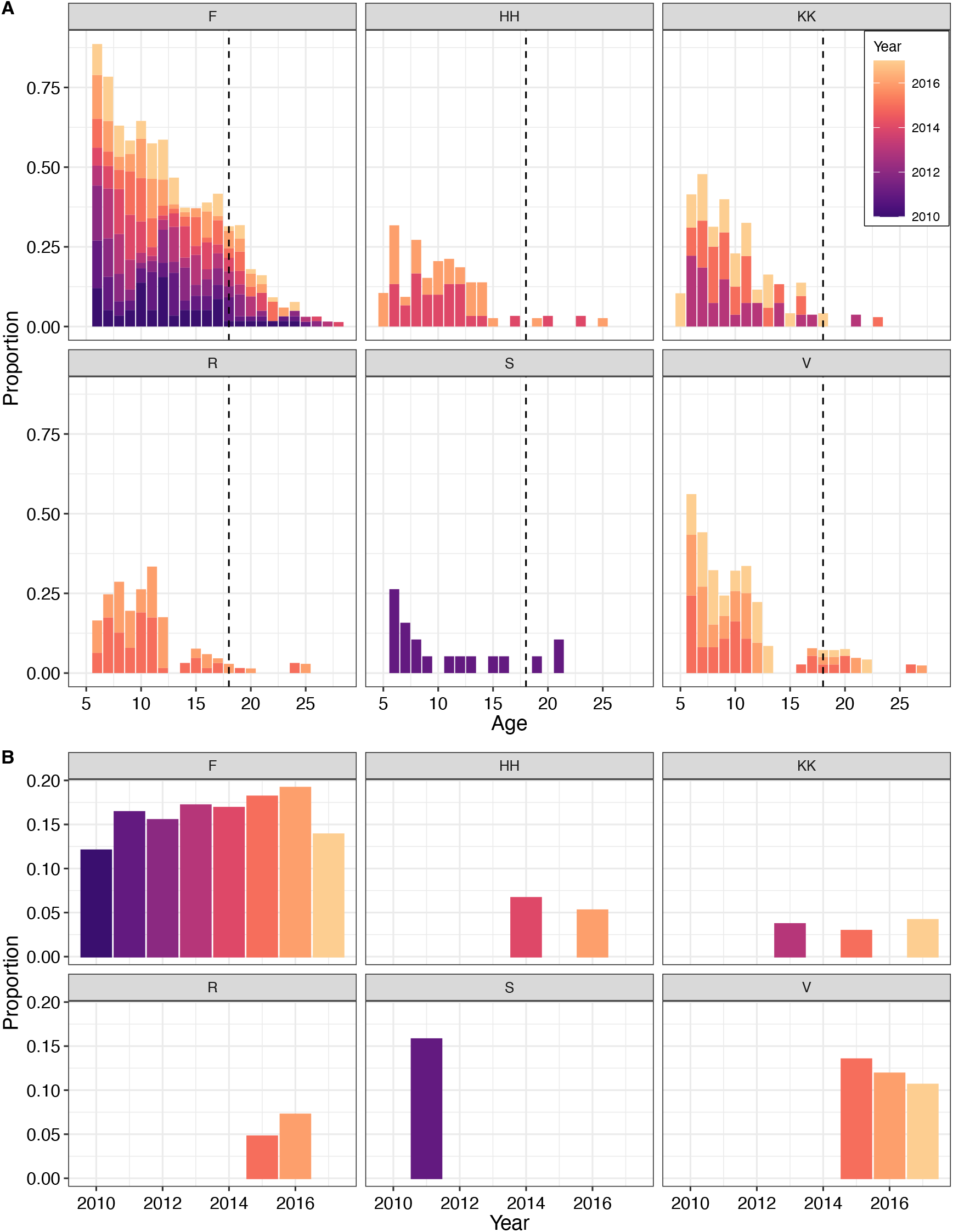
**(A)** Distribution of ages in each of the 6 empirical macaque groups for each year that those groups were observed (19 networks total observed between 2010-2017). The data are presented as a stacked bar chart, so to compare the proportion of x-year-olds in a single group across years one should compare the height of coloured bars within a single age. The dotted black line at age 18 indicates the median age of death in this population and the cut off at which we considered individuals “old” for the sake of calculating the proportion of old individuals in the group. See also Fig. S2. **(B)** Proportion of old individuals in each group for each year that group was observed. Years with no bars indicate years in which the group was not observed.

### Agent-based model of the relationship between age distribution and network structure

We expected that age-based changes in patterns of direct association would scale up to affect overall network structure. However, our empirical data revealed no effect of the proportion of old individuals in the network on global network metrics (see Results section below). To better understand these results, we built an individual-based model where we could manipulate age-based differences in sociality in isolation of other variables to better explore how different age distributions would be expected to affect overall network structure, and why such scaling up of age-based differences in social behaviour may not be detectable in our data. Our general approach was to simulate artificial populations that exhibited the social age-dependencies observed in the rhesus macaque system on Cayo Santiago and use these to investigate the relationship between age distribution and global network metrics. As mentioned above, adult female rhesus macaques change two aspects of their sociality with age: their number of social partners, and the proportion of partners that are kin [6]. These individual changes in sociality across adulthood lead to differences between young and old individuals in both the number of social connections and probability of connecting with kin [6].

The model therefore simulates social networks of varying age distributions, in which the probability that two nodes (i.e., individuals) have an edge (i.e., a grooming link) depends on the age of the individuals and their and kinship to each other. Individuals belonged to two age categories (as in the empirical analysis for global structure; old adults and young adults, hereafter “old” and “young”) and two kin categories (kin and non-kin). This gives 6 dyad types, which we denote by the age categories of the two individuals and their kinship status (e.g., old/old kin). We established the linking probability (i.e., the chance of having an edge in the social network) for each dyad type based on the mean proportion of dyads of that type that had an edge across the 19 empirical macaque grooming networks. The linking probability for dyads that were old individuals who were related to each other (i.e., old/old kin) was 0.33; for old/old non-kin dyads it was 0.02; for old/young kin dyads it was 0.37; for old/young non-kin dyads it was 0.05; for young/young kin dyads it was 0.27 and for young/young non-kin dyads it was 0.08. We fixed group size at 50 individuals, which approximates the mean number of adult females in real groups on Cayo Santiago (mean ± SE = 50.7 ± 3.9). Each simulated network had 10 clusters of individuals who were related to each other (i.e., kin groups) with 5 individuals in each. The number of individuals within each kin group mirrors the mean number of close adult female kin (relatedness coefficient ≥ 0.125) that adult female rhesus macaques have on Cayo Santiago (mean ± SE = 5.2 ± 0.87). In the model, kinship between pairs of individuals was determined by their kin group membership: individuals from the same kin group were classed as kin and individuals from different kin groups were classed as non-kin.

Each simulation round (i.e., construction and quantification of one network) proceeded as follows: We first randomly drew the number of old individuals in the group (n_old_) from a uniform distribution with a set range [0, 50]. We then randomly assigned all 50 group members to age-groups (n_old_ old individuals and n_young_= 50-n_old_ young individuals) and kin groups. This allowed the age structure of the groups to vary across the range from only young individuals (0% old) to only old individuals (100% old). We then constructed the social network by drawing links, where the chance of each dyad getting a link depended on their dyad type (linking probabilities given above). To determine whether a dyad was given a link, we extracted a random value from a binomial distribution with a sample of 1 and probability equal to the linking probability of the type of dyad. If the extracted value was 1, the dyad was given a link, if it was 0, the dyad was not given a link. A schematic representation of the various steps of the model-building process can be found in Fig S3.

To confirm that the model was working as intended, we ran 10000 simulations (i.e., we generated 10000 networks), from which we calculated the proportion of dyads of each type that had a link. We confirmed that the means of these proportions (old/old kin = 0.33; old/old non-kin = 0.02; old/young kin = 0.37; old/young non-kin = 0.05; young/young kin = 0.27; young/young non-kin = 0.08) corresponded well to their respective empirical linking probability (see above, and Fig. S4). We also confirmed that our model was not highly sensitive to our input linking probabilities (see Supplementary Methods, Table S1, Fig. S5 for more information). Given that our input values are only estimates of the real values, robustness to these values is important for drawing general conclusions about the potential effects in the real system.

To investigate the general relationship between age distribution and network structure, we ran 100000 replicates of the simulation (i.e., generated 100000 networks) where we allowed the proportion of old individuals in the network to range from 0.00 to 1.00 and visualised the relationship between the proportion of old individuals and network mean degree, diameter, and transitivity. Additionally, we used the agent-based model to investigate if we should expect to find relationships between the proportion of old individuals and network metrics in the empirical analysis. To do this, we limited the proportion of old individuals in each network to between 0.04 and 0.20 (mirroring the empirical variation in age distributions), and we ran 19 simulations (mirroring the 19 empirical networks). We repeated this 50 times (i.e., 50 sets of 19 networks) to gauge consistency of the results.

## Results

### Relationship between age and indirect measures of network connectedness

In line with our predictions, female macaques did not show any change in eigenvector centrality with age (within-individual age: *β* = -0.04; 95% CI = -0.30, 0.24; Fig. 2A-B, Table S2). That is, as females aged, the strength of their relationships to their partners, and to their partners’ partners, was stable. We did find evidence of a within-individual decline in betweenness with age. However, this effect was rank dependent and seems to be driven primarily by a decline in betweenness as high-ranking individuals got older (within-individual age:rankH: *β* = -0.35; 95% CI = -0.41, -0.29; Fig. 2C-D, Table S3), although mid-ranking individuals also showed greater declines in betweenness with age than did low-ranking females (within-individual age:rankM: *β* = -0.14; 95% CI = -0.18, -0.09; Fig. 2C-D, Table S3). Therefore, some, but not all individuals became less effective at reaching disparate nodes in the network as they got older. Individuals showed a reduction in their closeness with age (within-individual age: *β* = -0.18; 95% CI = -0.35, -0.02; Fig. 2E-F, Table S4), meaning they were harder to reach and be reached in the network, as we predicted. Contrary to our predictions, there was no within-individual effect of age on clustering coefficient (within-individual age: *β* = 0.04; 95% CI = -0.06, 0.14; Fig. 2G-H, Table S5). That is, individual cliquishness was stable as individuals aged.

**Figure 2.**
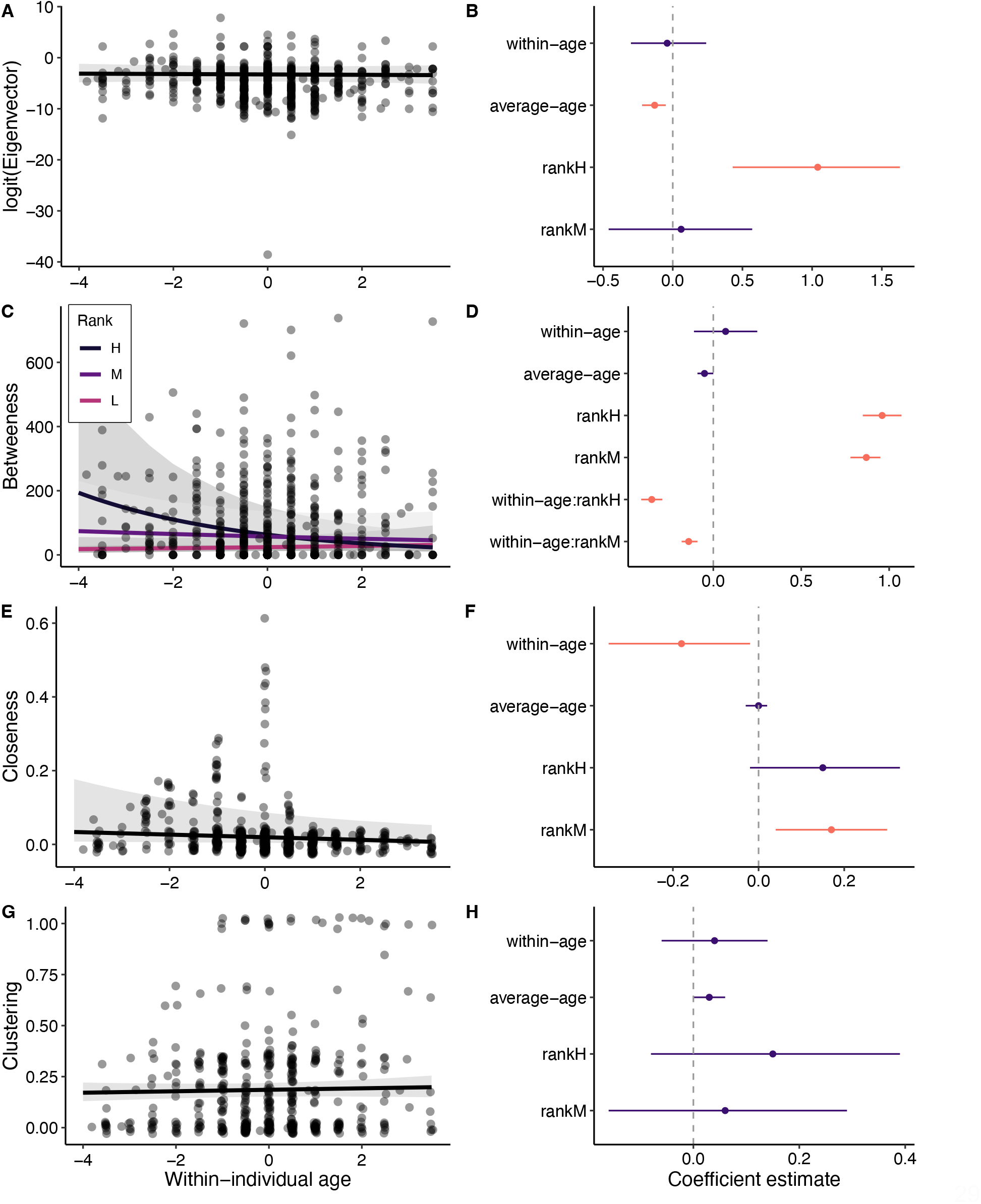
Relationship between within-individual changes in age and indirect measures of connectedness including (A & B) eigenvector centrality, (C & D) betweenness, (E & F) closeness, and (G & H) clustering coefficient in female rhesus macaques. In **A, C, E, G** the points represent raw data. Shaded grey bars indicate 95% confidence intervals around the predicted values. In **B, D, F, H** effect sizes and 95% credible intervals for all fixed effects and interaction terms are shown. Instances where the 95% CI overlaps zero are coloured in purple.

### Relationship between age distribution and network structure

#### Empirical results

Contrary to our predictions, we found no evidence that groups with a greater number of old individuals were structured differently from groups with fewer old individuals. There was no overall effect of the proportion of old individuals in the group on mean degree (*β* = -2.76; 95% CI = -10.24, 5.25; n = 19; Fig. 3A, Table S6), or network diameter (*β* = -0.12; 95% CI = -3.93, 3.67; n = 19; Fig. 3B, Table S7). Therefore, networks were not more sparsely connected as the proportion of older animals increased. Older networks were also not more clustered or cliquish, as measured by transitivity (*β* = 0.20; 95% CI = -0.29, 0.70; n = 19; Fig. 3C, Table S8).

**Figure 3.**
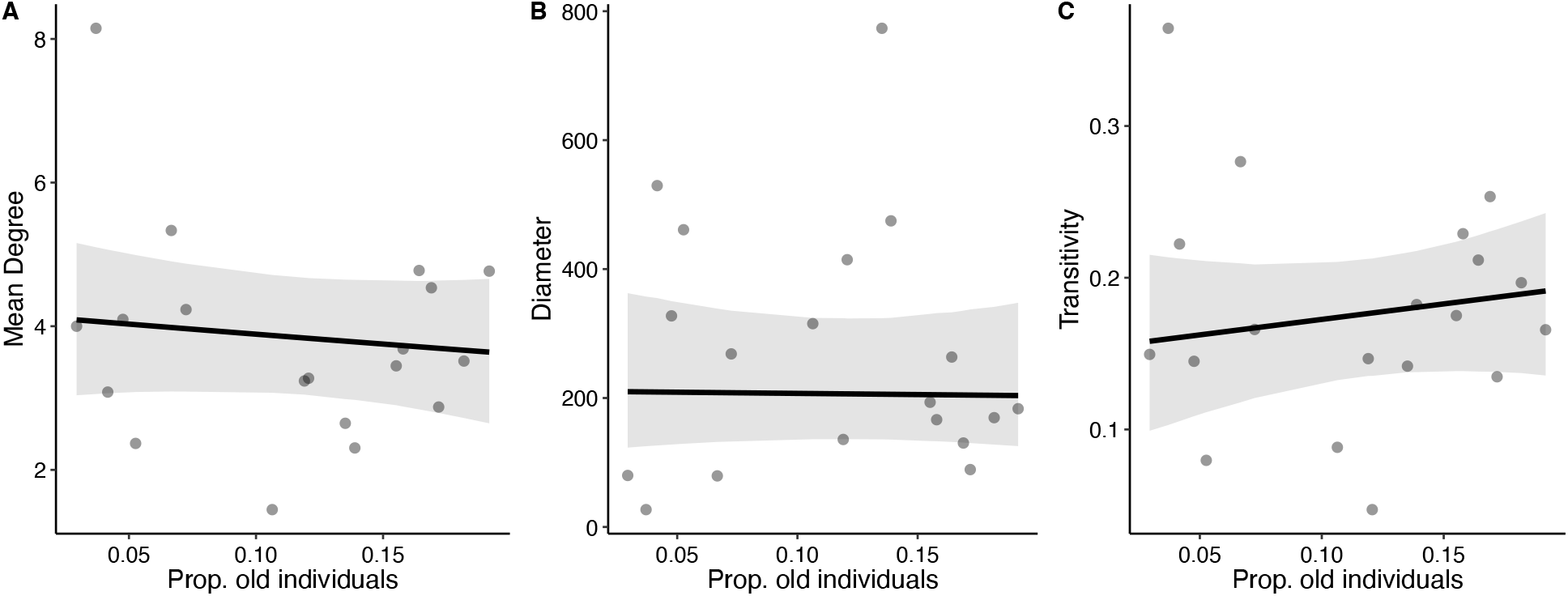
The relationship between the proportion of old individuals in female rhesus macaque networks and: (**A**) mean degree, (**B**) network diameter, and (**C**) network transitivity. Points represent raw data and shaded grey bars indicate 95% confidence intervals around the predicted values.

#### Agent-based model results

Different group demographics in age appeared to have important consequences for network structure in our simulations (Fig. 4; Fig. S6). Specifically, our 100,000 simulations where we allowed the proportion of old individuals in the network to range from 0.00 to 1.00 showed that, as the proportion of old individuals in the population increased, mean degree (i.e., the mean number of partners that each individual associated with) decreased (Fig. 4D), and the diameter of the network (i.e., the longest path length in the network) increased (Fig. 4E), both as expected. Network transitivity (i.e., the degree of clustering in the network; Fig. 4F) showed little variation in relation to age distribution.

**Figure 4.**
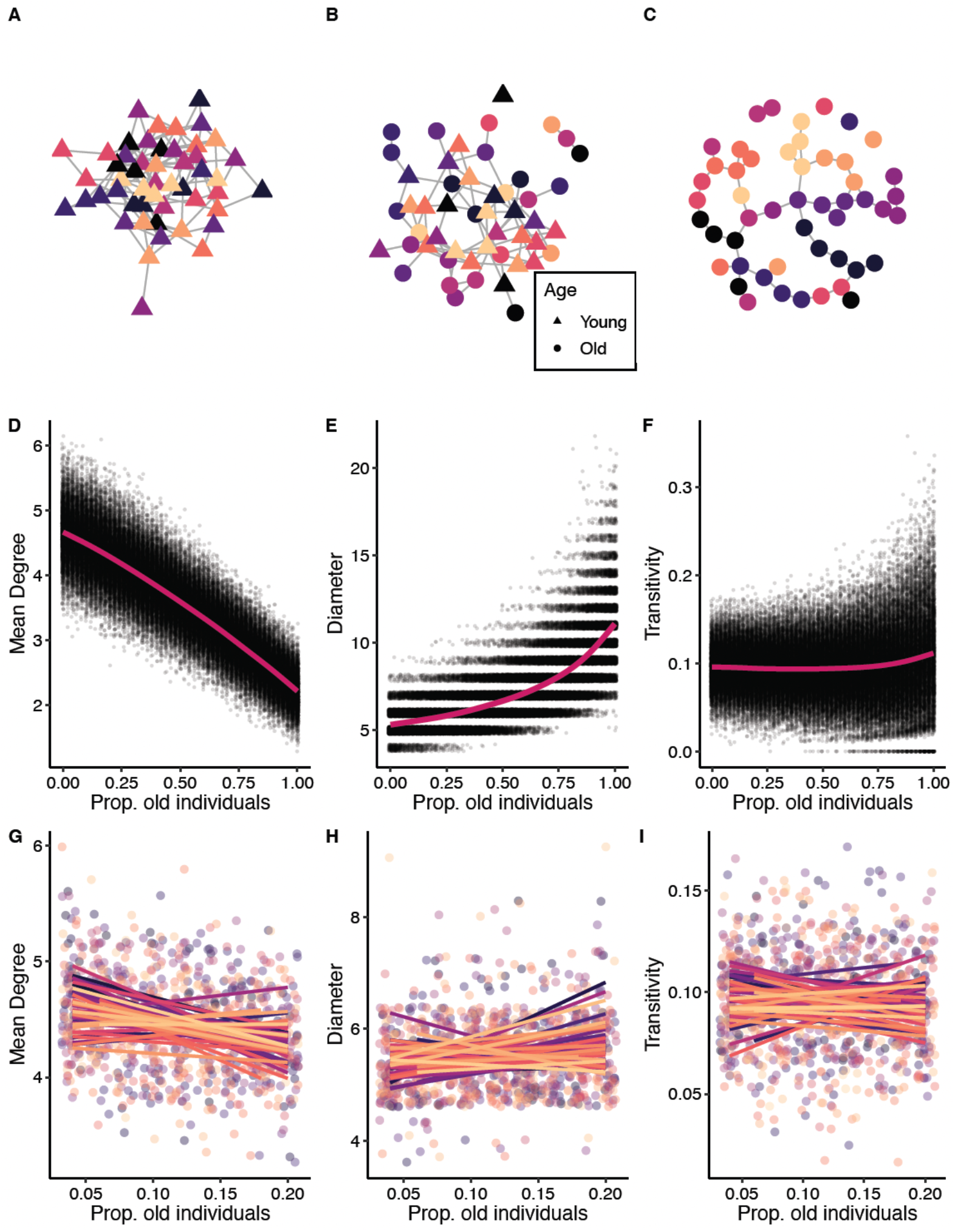
(**A-C**) Example networks from the agent-based model illustrating differences in network structure with the proportions of old individuals in the network equal to 0 (**A**), 0.5 (**B**) and 1 (**C**). Node colour represents kin groups where nodes of the same colour belong to the same kin group. (**D-F**) Results from the agent-based model showing relationship between the proportion of old individuals and (**D**) mean degree, (**E**) diameter, and (**F**) transitivity. Data points from the 100K simulations are shown with a small amount of jitter introduced to show overlapping points and are fitted with a smoothing term. (**G-I**) Relationship between the proportion of old individuals and (**G)** mean degree, (**H**) network diameter, and (**I**) transitivity based on 19 simulations from the agent-based model where the proportion of old individuals in the population was restricted between 0.04 and 0.20. We re-simulated these 19 networks with limited age variation 50 times to help ensure our results were robust. Each colour represents a different ‘bout’ of 19 networks. Data points are shown fitted with a linear smoothing term.

Importantly, these global metrics did not always exhibit linear relationships with network age (as might have initially been intuitively expected). The potential for non-linearity is important for understanding when the effects of age on network structure might be the most pronounced and most detectable in the real world. In particular, the increase in network diameter with an increasing proportion of old individuals showed an accelerating trend (Fig. 4E). That is, the relationship between network diameter and proportion of old individual was stronger (i.e., the slope was steeper) when the proportion of old individuals in the network was high, suggesting that effects of age on network structure might be most detectable when, for example, greater than 50% of the individuals in the network are old. While we did not find any clear change in transitivity with the proportion of old individuals in the network, there was also some evidence of an accelerating trend at the higher end of the age distribution. These metrics (transitivity, diameter) are driven to a greater extent by modularity in networks and the non-linear relationship could arise from the kin-biased nature of the relationships in our (modelled) study system. We also observed that transitivity and network diameter were more variable when there was a greater proportion of old individuals in the network. This suggests that changes in network structure with changing age distribution might be unpredictable, for example, networks might become more clustered or less clustered as the population gets older. Overall, these results suggest that compared to networks with more young individuals, networks with more old individuals are sparser and less cohesive, and can take on more varied structures.

When we limited the sample size and variation in age distribution to those of the empirical data (i.e., 19 networks where the proportion of old individuals was only allowed to vary between 0.04 and 0.20) we observed no clear change in network structure as the proportion of old individuals in the network increased (Fig. 4G-I).

## Discussion

Age has begun to emerge as an important attribute shaping the social decisions of individuals [6,7,14]. This implies that age is a potentially significant feature underlying the behaviour of animals in collectives. Here we have shown that age-based changes in social behaviour are likely to have important consequences for an individual’s position within the wider social network and may scale up to influence network topology. As female rhesus macaques aged, they showed declines in some, but not all of their indirect measures of connectedness. But despite age-based changes in both direct connectedness [6] and indirect connectedness (this paper) we detected no effect of age distribution on the overall structure of rhesus macaque networks. The agent-based model gives insight into this surprising result, as it implies that age-based differences in social behaviour do necessarily, as expected, scale up to affect network structure, but that these effects may be nonlinear and so may not always be easily detectable.

In line with our predictions, we found no evidence for changes in eigenvector centrality with age. This was expected because, despite reducing their number of social partners, female rhesus macaques maintain the amount of time they engage in social interactions as they get older [6], allowing them to continue to have strong connections to some of their partners, and (indirectly) to the partners of their partners. A similar pattern has been observed in a population of wild rhesus macaques in China whereby older individuals had fewer partners but exhibited similar weighted eigenvector centrality to those of younger individuals [34]. Unexpectedly, we did not find any evidence that clustering coefficient increased as individuals aged in this population. Females increase their preference for grooming kin as they age [6], suggesting that within-group (e.g., homophilic) tendencies increase with age, which should lead to greater clustering as females from the same matriline would cluster together in the network. However, although kin-biases increase with age, they exist for females of all ages [58]. It is therefore possible that ties with kin are the main driver of clustering coefficient scores regardless of age. The loss or removal of a few non-kin relationships as an individual ages could thus have a relatively minor impact on their clustering coefficient, leading to stable values for this metric across the adult lifespan. In general, the stability of weighted eigenvector centrality and clustering coefficient across the adult lifespan suggest that older animals can remain socially central and well-integrated in some respects, despite other aspects of their social life changing (e.g., fewer partners overall, and the patterns observed below). Old individuals may therefore continue to reap some of the advantages of social relationships [5].

As predicted, we found declines in betweenness and closeness with age, although some of these results were rank dependent. As individuals got older, particularly those that were higher ranking had less influence on and became less well-connected to the wider network.

Betweenness and closeness are associated with an individual’s level of reassociation tendency and within-group association [51]. In other words, the more likely individuals are to reassociate with the same partners, and to select partners of similar characteristics to themselves, the less they mix with the wider network and the less likely they are to connect distinct subgroups (i.e., have high betweenness) or be easily reached by all others in the network (i.e., have high closeness). We’ve previously shown that ageing rhesus macaque females have fewer partners, but engage with those partners more often, and increase their preference for close relatives. In other words, while mean dyadic association strength to associates increases with age, mixing with the wider network should decline [6]. Our betweenness and closeness results seem to reflect these underlying changes in behavioural patterns. The declines in betweenness with age were rank dependent, with mid-ranking and high-ranking rhesus macaque females showing a greater decline in betweenness with age than low-ranking individuals. This may simply be due to a floor effect – because low ranking individuals have such low betweenness to begin with, there is little capacity for further decline with age.

Generally, our findings demonstrate that how indirectly connected an individual is to their social world can change across their lifespan. How an individual is positioned in the wider social environment can modulate their exposure to information [26,31], parasites [18,77], and pathogens [20–22]. Individuals with high betweenness and closeness occupy a critical position in the acquisition and transfer of “goods” within a network [17,54,78]. Decreases in both of these measures of indirect connectedness with age may benefit aged individuals who may experience greater susceptibility to disease or illness as a result of immunosenescence [79]. Such benefits need not imply that individuals actively change complex network positions with age. By changing simple behavioural rules or processes with age, changes in polyadic ties with age are likely to emerge [51]. Measures of indirect connectedness have been linked to fitness proxies including future social status [80,81], survival [82,83], and reproductive success [32,64,81]. Indirect network metrics can be an even stronger predictor of fitness proxies [17,32] than measures of direct connections and may more strongly reflect the underlying behavioural rules that give rise to individual differences in sociality [51]. As such, documenting how ageing shapes the polyadic social world may be particularly relevant for understanding how changes in sociality across the lifespan influence patterns of senescence and fitness in later life.

While age is clearly associated with changes in the behaviour and social connectedness of animals living in groups [10,13,14], including in this population of rhesus macaques [6,60], we did not find any empirical evidence that age impacts the overall structure of those groups. We found no relationship between age distribution and network mean degree, diameter, or transitivity in the observed macaque data. This was surprising to us given that we expected that age-based changes at the individual level would scale up to the network level. That is, it should be self-evident that a network with more old individuals who each interact with fewer partners would, for example, have a lower mean degree. Given that we did not find empirical evidence of this, we used an agent-based modelling approach to try to understand why these results might emerge and whether under a scenario where everything else is equal, we could recreate these expected effects on network structure.

By modelling age-based differences in two interaction patterns (number of social partners and the tendency to link with kin) based on the age-based changes we observed in the female macaques [6], we found that the age composition of a group can have important consequences for its cohesiveness and connectedness. As predicted, mean degree declined and diameter increased as the proportion of old individuals increased, while transitivity exhibited no strong relationship to age-distribution. Interestingly, these effects did not necessarily scale in the linear manner that might be expected in response to a linear increase in proportion of old individuals in the network. Network diameter showed a steeper increase as networks became older, suggesting that the strength of the effect, and thereby the ability to statistically detect an effect in a real-world case, might depend on where along the continuum of age distributions one’s data lie. The combination of this and a limited sample size could potentially have led to the null result in our empirical analysis. This interpretation is supported by the fact that when we limited the age variation and sample size (number of networks) of our model to match that of the empirical data (i.e., only ran the model for 19 simulations and allowed the proportion of old individuals in the network to range between 0.4-0.20) there was no clear relationship between the proportion of old individuals and network structure.

The results of our model suggest that populations made up of very large numbers of young, or old, individuals may have detectable levels of divergence in network structure, whereas populations within a smaller range of age distributions may not. Sample size also appears to have played an important role in our ability to detect an effect of network age on network structure. Each “bout” of 19 networks (simulations) produced a slightly different result. Sometimes slopes were positive, sometimes negative, sometimes flat, suggesting there is a fair amount of stochasticity in our ability to detect the true effect when the sample size is small, even in the model world where there are no other confounding factors. While nineteen rhesus macaque networks from 6 groups collected across 8 years represents a considerable investment of research time and effort, it may not have been enough to detect a clear pattern. However, it is also possible that our empirical results reflect a true null result if there are processes occurring that our model has not accounted for. Our model is a simplistic version of the real world and does not fully replicate the complete suite of changes that might occur as individuals age. For example, our model does not allow for potential rewiring between nodes which might occur if younger individuals choose to build new ties in response to the loss of ties with older individuals. Such rewiring could explain why some measures of network structure do not change despite individual declines in sociality with age.

While the general result from the agent-based model was expected, these simulations help to inform our understanding of the empirical data and offer us a glimpse into how age-based differences in sociality may affect network structure and when these effects might be detected. The fact that, even in our simplified model where there is nothing to obscure individual linking probabilities from scaling up to affect global network metrics, changes in some measures of network structure appear most likely to be detected in very old populations raises the question of when such effects are likely to be detectable in wild systems. The macaques of Cayo Santiago are a managed population that are food supplemented and have no predators. As such the proportion of old individuals seems likely to be higher than in many wild populations and aligns with other natural but non-predated systems (e.g., red deer [84]) where the proportion of old individuals in the population appears to top out at about 20%. Our model suggests that the effects of age on network structure are also present at this lower end of the age-distribution, but are less strong, or negligible, for some network measures. Hence, detecting an effect in real systems could necessitate large amounts of data. We note, however, that the effect on the network measures (the shapes of the curves) will depend both on how sociality changes with age, and the social organisation of the system (such as relatedness patterns), and while other systems may be similar to the one here investigated in these regards, making predictions for detectability of social ageing effects in general would require more comprehensive modelling.

Future work in other systems, conditions, and contexts could reveal consequences of age-based changes in sociality for network structure that are even more stark. For example, greater differences between age classes in the overall probability of forming a social tie may be expected in societies that experience selective disappearance of more social individuals or where declines in sociality with age are exacerbated by partners dying and not being replaced (which we do not observe in this system [6]). The kin structure of populations is also likely to be important if the probability of connecting decreases with age but there is no compensatory increase in the probability of forming ties with kin. This could lead to stronger absolute effects on global network metrics, including substantial changes in transitivity, and thus would likely have further consequences for network connectivity and spreading processes. While our analyses bin individuals into age categories of “old” and “young”, there may be much more nuanced effects of age on network structure if the full age distribution of a population or group is taken into account.

It is also important to recognize that the results presented here only reflect how changes in female sociality across adulthood are expected to influence adult female social networks. Understanding how social networks of entire groups are affected by age-based changes in behaviour, more broadly, will necessitate incorporating an understanding of the patterns of social ageing in both sexes, as well as an understanding of how interactions with juveniles shift as adults get older. While much less is known about the social ageing trajectories of males, some studies suggest that they show lower levels of connectedness at older ages, similar to those observed in females [13,33]. But even in cases where social ageing trajectories are similar across the sexes, changes in socio-sexual interactions with age in addition to shifting interactions between age classes are likely to produce much more complex network dynamics than observed here, necessitating further research.

Despite these limitations, our study provides a first glimpse into the effects that age-based changes in behaviour might have on network structure, offering a deeper understanding of the potential importance of demographic changes for the structure and function of collectives. Much attention has been given to the potential for the social structure of populations to be affected by the removal of older individuals, for example through practices like trophy hunting or fishing [85,86]. This is relevant given that older individuals can play a critical role in their social groups through leadership [4,39], intergenerational transfers [40], and the stabilisation of social relationships [87]. Despite the importance of the loss of older individuals from networks, our results suggest that asking what happens when populations contain large numbers of older individuals may be an equally salient question.

In humans, population ageing is poised to be one of the most significant social transformations of the 21^st^ century, with the global number of older persons projected to more than double over the next three decades [88], facilitated by falling fertility rates and increasing longevity [89,90]. Meanwhile, in other group-living species, anthropogenic challenges, including climate change, poaching, disease outbreaks, and pollution, among others, are likely to cause major population declines and demographic shifts [91,92]. The consequences of declining fertility and/or increasing mortality rates may be further exacerbated by the concomitant shifts in age structure and resulting implications for network structure and function. For instance, declines in cohesiveness and structural connectedness in older networks may limit or slow the transfer of information [24,93], restrict the potential for cooperation [94] or reduce the stability of populations [30], feeding back to further limit population growth. Alternatively, such changes could reduce the vulnerability of populations to infectious diseases or parasites [30,95] or enhance cooperation if network modularity is increased [96]. Consequently, through its effects on network structure, shifting age demography may have broader implications than previously appreciated for group dynamics and persistence and warrants further research.

## Acknowledgments

Thank you to Kate Laskowski and Christos Ioannou for organising a stimulating meeting and to the Royal Society for hosting. We are very grateful to K. Laskowski, J. Firth and two anonymous reviewers for constructive feedback on a previous version of this manuscript. We thank the Caribbean Primate Research Center for maintaining the Cayo Santiago population and for access to the study site, and all the field technicians who have contributed to the long-term behavioural database over the years. We are grateful to members of the Centre for Research in Animal Behaviour at the University of Exeter, as well as Social Network Club, for thoughtful discussion during the manuscript’s development. This work was supported by the following grants from the National Institute of Health (NIH): grant nos R01-AG060931, R00-AG051764, R01-MH096875, R37-MH109728, R01-MH108627, R01-MH118203, U01MH121260, R01-NS123054 and the Kaufman Foundation: grant no KA2019-105548. The Cayo Santiago Field Station is supported by the Office of Research Infrastructure Programs of the NIH (2P40OD012217).

## Data availability

All data and code associated with the analyses will be made publicly available on the Figshare Repository following acceptance of the manuscript.

## References

1. Lucas ER, Keller L. 2020 The co-evolution of longevity and social life. Funct. Ecol. 34, 76–87.

2. Johnstone RA, Cant MA. 2010 The evolution of menopause in cetaceans and humans: the role of demography. Proc. Biol. Sci. 277, 3765–3771.

3. Croft DP, Brent LJN, Franks DW, Cant MA. 2015 The evolution of prolonged life after reproduction. Trends Ecol. Evol. 30, 407–416.

4. Brent LJN, Franks DW, Foster EA, Balcomb KC, Cant MA, Croft DP. 2015 Ecological knowledge, leadership, and the evolution of menopause in killer whales. Curr. Biol. 25, 746–750.

5. Snyder-Mackler N et al. 2020 Social determinants of health and survival in humans and other animals. Science 368. (doi:10.1126/science.aax9553)

6. Siracusa ER, Negron-Del Valle JE, Phillips D, Platt ML, Higham JP, Snyder-Mackler N, Brent LJN. 2022 Within-individual changes reveal increasing social selectivity with age in rhesus macaques. bioRxiv., 2022.05.31.494118. (doi:10.1101/2022.05.31.494118)

7. Machanda ZP, Rosati AG. 2020 Shifting sociality during primate ageing. Philosophical Transactions of the Royal Society B: Biological Sciences. 375, 20190620. (doi:10.1098/rstb.2019.0620)

8. Weiss MN et al. 2021 Age and sex influence social interactions, but not associations, within a killer whale pod. Proceedings of the Royal Society B: Biological Sciences 288, 20210617.

9. Kroeger SB, Blumstein DT, Martin JGA. 2021 How social behaviour and life-history traits change with age and in the year prior to death in female yellow-bellied marmots. Philos. Trans. R. Soc. Lond. B Biol. Sci. 376, 20190745.

10. Albery GF, Clutton-Brock TH, Morris A, Morris S. 2021 Ageing red deer alter their spatial behaviour and become less social. bioRxiv

11. Verhulst S, Geerdink M, Salomons HM, Boonekamp JJ. 2014 Social life histories: jackdaw dominance increases with age, terminally declines and shortens lifespan. Proc. Biol. Sci. 281, 20141045.

12. Wrzus C, Hänel M, Wagner J, Neyer FJ. 2013 Social network changes and life events across the life span: a meta-analysis. Psychol. Bull. 139, 53–80.

13. Rosati AG, Hagberg L, Enigk DK, Otali E, Emery Thompson M, Muller MN, Wrangham RW, Machanda ZP. 2020 Social selectivity in aging wild chimpanzees. Science 370, 473–476.

14. Almeling L, Hammerschmidt K, Sennhenn-Reulen H, Freund AM, Fischer J. 2016 Motivational Shifts in Aging Monkeys and the Origins of Social Selectivity. Curr. Biol. 26, 1744–1749.

15. Schino G, Pinzaglia M. 2018 Age-related changes in the social behavior of tufted capuchin monkeys. Am. J. Primatol. 80, e22746.

16. Hinde RA. 1976 Interactions, Relationships and Social Structure. Man 11, 1–17.

17. Brent LJN. 2015 Friends of friends: are indirect connections in social networks important to animal behaviour? Anim. Behav. 103, 211–222.

18. White LA, Forester JD, Craft ME. 2017 Using contact networks to explore mechanisms of parasite transmission in wildlife. Biol. Rev. Camb. Philos. Soc. 92, 389–409.

19. Firth JA. 2020 Considering Complexity: Animal Social Networks and Behavioural Contagions. Trends Ecol. Evol. 35, 100–104.

20. Drewe JA. 2010 Who infects whom? Social networks and tuberculosis transmission in wild meerkats. Proc. Biol. Sci. 277, 633–642.

21. Morrison RE, Mushimiyimana Y, Stoinski TS, Eckardt W. 2021 Rapid transmission of respiratory infections within but not between mountain gorilla groups. Sci. Rep. 11, 19622.

22. Weber N, Carter SP, Dall SRX, Delahay RJ, McDonald JL, Bearhop S, McDonald RA. 2013 Badger social networks correlate with tuberculosis infection. Curr. Biol. 23, R915–6.

23. Godfrey SS. 2013 Networks and the ecology of parasite transmission: A framework for wildlife parasitology. Int. J. Parasitol. Parasites Wildl. 2, 235–245.

24. Aplin LM, Farine DR, Morand-Ferron J, Cockburn A, Thornton A, Sheldon BC. 2015 Experimentally induced innovations lead to persistent culture via conformity in wild birds. Nature 518, 538–541.

25. Evans JC, Silk MJ, Boogert NJ, Hodgson DJ. 2020 Infected or informed? Social structure and the simultaneous transmission of information and infectious disease. Oikos 129, 1271–1288.

26. Claidière N, Messer EJE, Hoppitt W, Whiten A. 2013 Diffusion Dynamics of Socially Learned Foraging Techniques in Squirrel Monkeys. Curr. Biol. 23, 1251–1255.

27. King AJ, Sueur C, Huchard E, Cowlishaw G. 2011 A rule-of-thumb based on social affiliation explains collective movements in desert baboons. Anim. Behav. 82, 1337–1345.

28. Bode NWF, Wood AJ, Franks DW. 2011 The impact of social networks on animal collective motion. Anim. Behav. 82, 29–38.

29. Cantor M et al. 2021 The importance of individual-to-society feedbacks in animal ecology and evolution. J. Anim. Ecol. 90, 27–44.

30. Kurvers RHJM, Krause J, Croft DP, Wilson ADM, Wolf M. 2014 The evolutionary and ecological consequences of animal social networks: emerging issues. Trends Ecol. Evol. 29, 326–335.

31. Aplin LM, Farine DR, Morand-Ferron J, Sheldon BC. 2012 Social networks predict patch discovery in a wild population of songbirds. Proc. Biol. Sci. 279, 4199–4205.

32. Cheney DL, Silk JB, Seyfarth RM. 2016 Network connections, dyadic bonds and fitness in wild female baboons. R Soc Open Sci 3, 160255.

33. Rathke E-M, Fischer J. 2021 Social aging in male and female Barbary macaques. Am. J. Primatol., e23272.

34. Liao Z, Sosa S, Wu C, Zhang P. 2018 The influence of age on wild rhesus macaques” affiliative social interactions. Am. J. Primatol. 80. (doi:10.1002/ajp.22733)

35. Brent LJN, Ruiz-Lambides A, Platt ML. 2017 Persistent social isolation reflects identity and social context but not maternal effects or early environment. Scientific Reports. 7. (doi:10.1038/s41598-017-18104-4)

36. Wey TW, Blumstein DT. 2010 Social cohesion in yellow-bellied marmots is established through age and kin structuring. Anim. Behav. 79, 1343–1352.

37. Nussey DH, Coulson T, Festa-Bianchet M, Gaillard J-M. 2008 Measuring Senescence in Wild Animal Populations: Towards a Longitudinal Approach. Funct. Ecol. 22, 393–406.

38. Siracusa ER, Higham JP, Snyder-Mackler N, Brent LJN. 2022 Social ageing: exploring the drivers of late-life changes in social behaviour in mammals. Biol. Lett. 18, 20210643.

39. Allen CRB, Brent LJN, Motsentwa T, Weiss MN, Croft DP. 2020 Importance of old bulls: leaders and followers in collective movements of all-male groups in African savannah elephants (Loxodonta africana). Scientific Reports. 10. (doi:10.1038/s41598-020-70682-y)

40. McComb K, Moss C, Durant SM, Baker L, Sayialel S. 2001 Matriarchs as repositories of social knowledge in African elephants. Science 292, 491–494.

41. Mueller T, O”Hara RB, Converse SJ, Urbanek RP, Fagan WF. 2013 Social learning of migratory performance. Science 341, 999–1002.

42. Lee HC, Teichroeb JA. 2016 Partially shared consensus decision making and distributed leadership in vervet monkeys: older females lead the group to forage. Am. J. Phys. Anthropol. 161, 580–590.

43. Allen CRB, Croft DP, Brent LJN. 2021 Reduced older male presence linked to increased rates of aggression to non-conspecific targets in male elephants. Proc. Biol. Sci. 288, 20211374.

44. Slotow R, van Dyk G, Poole J, Page B, Klocke A. 2000 Older bull elephants control young males. Nature 408, 425–426.

45. Bourjade M, des Roches A de B, Hausberger M. 2009 Adult-Young Ratio, a Major Factor Regulating Social Behaviour of Young: A Horse Study. PLoS ONE. 4, e4888. (doi:10.1371/journal.pone.0004888)

46. Williams R, Lusseau D. 2006 A killer whale social network is vulnerable to targeted removals. Biol. Lett. 2, 497–500.

47. Cook PA, Baker OM, Costello RA, Formica VA, Brodie ED 3rd. 2022 Group composition of individual personalities alters social network structure in experimental populations of forked fungus beetles. Biol. Lett. 18, 20210509.

48. Shizuka D, Johnson AE. 2019 How demographic processes shape animal social networks. Behav. Ecol. 31, 1–11.

49. Pinter-Wollman N. 2015 Persistent variation in spatial behavior affects the structure and function of interaction networks. Curr. Zool. 61, 98–106.

50. Ilany A, Akçay E. 2016 Social inheritance can explain the structure of animal social networks. Nat. Commun. 7, 12084.

51. Firth JA, Sheldon BC, Brent LJN. 2017 Indirectly connected: simple social differences can explain the causes and apparent consequences of complex social network positions. Proc. Biol. Sci. 284. (doi:10.1098/rspb.2017.1939)

52. Ilany A. 2019 Complex societies, simple processes: a comment on Shizuka and Johnson. Behav. Ecol. 31, 13–13.

53. Chiou KL et al. 2020 Rhesus macaques as a tractable physiological model of human ageing. Philos. Trans. R. Soc. Lond. B Biol. Sci. 375, 20190612.

54. Wey T, Blumstein DT, Shen W, Jordán F. 2008 Social network analysis of animal behaviour: a promising tool for the study of sociality. Anim. Behav. 75, 333–344.

55. Widdig A et al. 2016 Genetic studies on the Cayo Santiago rhesus macaques: A review of 40 years of research. Am. J. Primatol. 78, 44–62.

56. Pavez-Fox MA et al. 2022 Reduced injury risk links sociality to survival in a group-living primate. bioRxiv., 2022.04.05.487140. (doi:10.1101/2022.04.05.487140)

57. Bernstein IS, Judge PG, Ruehlmann TE. 1993 Kinship, association, and social relationships in rhesus monkeys (Macaca mulatta). Am. J. Primatol. 31, 41–53.

58. Berman CM. 2016 Primate Kinship: Contributions from Cayo Santiago. Am. J. Primatol. 78, 63–77.

59. Blomquist GE, Sade DS, Berard JD. 2011 Rank-Related Fitness Differences and Their Demographic Pathways in Semi-Free-Ranging Rhesus Macaques (Macaca mulatta). Int. J. Primatol. 32, 193–208.

60. Brent LJN, Ruiz-Lambides A, Platt ML. 2017 Family network size and survival across the lifespan of female macaques. Proc. Biol. Sci. 284. (doi:10.1098/rspb.2017.0515)

61. Datta S. 1988 The acquisition of dominance among free-ranging rhesus monkey siblings. Anim. Behav. 36, 754–772.

62. Rosati AG, Santos LR. 2017 Tolerant Barbary macaques maintain juvenile levels of social attention in old age, but despotic rhesus macaques do not. Anim. Behav. 130, 199–207.

63. Altmann J. 1974 Observational study of behavior: sampling methods. Behaviour 49, 227–267.

64. Brent LJN, Heilbronner SR, Horvath JE, Gonzalez-Martinez J, Ruiz-Lambides A, Robinson AG, Skene JHP, Platt ML. 2013 Genetic origins of social networks in rhesus macaques. Sci. Rep. 3, 1042.

65. Ellis S, Snyder-Mackler N, Ruiz-Lambides A, Platt ML, Brent LJN. 2019 Deconstructing sociality: the types of social connections that predict longevity in a group-living primate. Proc. Biol. Sci. 286, 20191991.

66. Whitehead H. 2008 Analyzing Animal Societies. University of Chicago Press.

67. R Core Team. 2021 R: A language and environment for statistical computing. Vienna, Austria: R Foundation for Statistical Computing. https://www.R-project.org/.

68. Csardi G, Nepusz T. 2006 The igraph software package for complex network research. InterJournal, Complex Systems, 1695. https://igraph.org.

69. Stan Development Team. 2021 Stan Modeling Language Users Guide and Reference Manual, 2.29. https://mc-stan.org.

70. Bürkner PC. 2017 brms: An R package for Bayesian multilevel models using Stan. Journal of Statistical Software 80. doi:10.18637/jss.v080.i01.

71. Hoffman CL, Higham JP, Mas-Rivera A, Ayala JE, Maestripieri D. 2010 Terminal investment and senescence in rhesus macaques (Macaca mulatta) on Cayo Santiago. Behav. Ecol. 21, 972–978.

72. Schwartz SM, Kemnitz JW. 1992 Age- and gender-related changes in body size, adiposity, and endocrine and metabolic parameters in free-ranging rhesus macaques. Am. J. Phys. Anthropol. 89, 109–121.

73. Hope GM, Dawson WW, Engel HM, Ulshafer RJ, Kessler MJ, Sherwood MB. 1992 A primate model for age related macular drusen. Br. J. Ophthalmol. 76, 11–16.

74. DeRousseau CJ. 1985 Aging in the musculoskeletal system of rhesus monkeys: III. Bone loss. Am. J. Phys. Anthropol. 68, 157–167.

75. Sanchez Rosado MR et al. 2021 Sociodemographic effects on immune cell composition in a free-ranging non-human primate. bioRxiv., 2021.12.06.471383. (doi:10.1101/2021.12.06.471383)

76. Van de Pol M, Wright J. 2009 A simple method for distinguishing within-versus between-subject effects using mixed models. Anim. Behav. 77, 753.

77. Rimbach R, Bisanzio D, Galvis N, Link A, Di Fiore A, Gillespie TR. 2015 Brown spider monkeys (Ateles hybridus): a model for differentiating the role of social networks and physical contact on parasite transmission dynamics. Philos. Trans. R. Soc. Lond. B Biol. Sci. 370. (doi:10.1098/rstb.2014.0110)

78. Freeman LC. 1977 A Set of Measures of Centrality Based on Betweenness. Sociometry 40, 35–41.

79. Peters A, Delhey K, Nakagawa S, Aulsebrook A, Verhulst S. 2019 Immunosenescence in wild animals: meta-analysis and outlook. Ecol. Lett. 22, 1709–1722.

80. McDonald DB. 2007 Predicting fate from early connectivity in a social network. Proc. Natl. Acad. Sci. U. S. A. 104, 10910–10914.

81. Gilby IC, Brent LJN, Wroblewski EE, Rudicell RS, Hahn BH, Goodall J, Pusey AE. 2013 FITNESS BENEFITS OF COALITIONARY AGGRESSION IN MALE CHIMPANZEES. Behav. Ecol. Sociobiol. 67, 373–381.

82. Stanton MA, Mann J. 2012 Early social networks predict survival in wild bottlenose dolphins. PLoS One 7, e47508.

83. Lehmann J, Majolo B, McFarland R. 2016 The effects of social network position on the survival of wild Barbary macaques, Macaca sylvanus. Behavioral Ecology. 27, 20–28. (doi:10.1093/beheco/arv169)

84. Clutton-Brock TH, Coulson T. 2002 Comparative ungulate dynamics: the devil is in the detail. Philos. Trans. R. Soc. Lond. B Biol. Sci. 357, 1285–1298.

85. Birkeland C, Dayton PK. 2005 The importance in fishery management of leaving the big ones. Trends Ecol. Evol. 20, 356–358.

86. Roach DA, Carey JR. 2014 Population biology of aging in the wild. Annu. Rev. Ecol. Evol. Syst. 45, 421–443.

87. Archie EA, Chiyo PI. 2012 Elephant behaviour and conservation: social relationships, the effects of poaching, and genetic tools for management. Mol. Ecol. 21, 765–778.

88. Nations U. 2019 World Population Ageing 2019 Highlights. pdf.

89. Oeppen J, Vaupel JW. 2002 Demography. Broken limits to life expectancy. Science 296, 1029–1031.

90. Harper S. 2014 Economic and social implications of aging societies. Science 346, 587–591.

91. Selwood KE, McGeoch MA, Mac Nally R. 2015 The effects of climate change and land-use change on demographic rates and population viability. Biol. Rev. Camb. Philos. Soc. 90, 837–853.

92. Moss CJ. 2001 The demography of an African elephant (Loxodonta africana) population in Amboseli, Kenya. J. Zool. 255, 145–156.

93. Cantor M, Whitehead H. 2013 The interplay between social networks and culture: theoretically and among whales and dolphins. Philos. Trans. R. Soc. Lond. B Biol. Sci. 368, 20120340.

94. Croft DP, James R, Thomas POR, Hathaway C, Mawdsley D, Laland KN, Krause J. 2006 Social structure and co-operative interactions in a wild population of guppies (Poecilia reticulata). Behav. Ecol. Sociobiol. 59, 644–650.

95. VanderWaal KL, Atwill ER, Hooper S, Buckle K, McCowan B. 2013 Network structure and prevalence of Cryptosporidium in Belding”s ground squirrels. Behav. Ecol. Sociobiol. 67, 1951–1959.

96. Marcoux M, Lusseau D. 2013 Network modularity promotes cooperation. J. Theor. Biol. 324, 103–108.

