## Supplementary Materials for "Ageing in a collective: The impact of ageing individuals on social network structure"

**This PDF file includes:**

Supplementary Methods
Tables S1 to S8
Figures S1 to S6

**Supplementary Methods**

To ensure that the conclusions from the model were not highly sensitive to the linking probability estimates, we investigated how the shape of the relationships between the proportion of old individuals and the network metrics changed when the difference between linking probabilities of old and young individuals became smaller. Our endpoint was the case where there was no difference between the social behaviour of old and young individuals (i.e., their respective linking probabilities converged to the same values). We set the linking probabilities to converge at 0.3 for kin dyads and 0.05 for non-kin dyads (Table S1) because those values represent mid-points between the linking probabilities of old/old (0.330) and young/young (0.270) kin dyads, and old/old (0.02) and young/young (0.08) non-kin dyads. Our goal was to remove the difference between the two age groups in incremental steps while keeping everything else equal, including an effect of kinship on linking. Therefore, the convergence points were different for kin and non-kin dyads.

We ran the model for this extreme case, for the original data-based values, and for five other sets of linking probabilities, which were at regular distances between the original values and the endpoint values (Table S1). For each of these cases, we ran 10000 simulations. Under high sensitivity, any observed correlations would disappear quickly (e.g., at the first reduction of difference in linking probabilities between age groups), whereas a more gradual reduction would indicate robustness to the input. The effects of the proportion of old individuals on network structure were only gradually reduced when the difference in social behaviour between young and old individuals was reduced, indicating that our conclusions are robust to the input parameter values (Fig. S5).

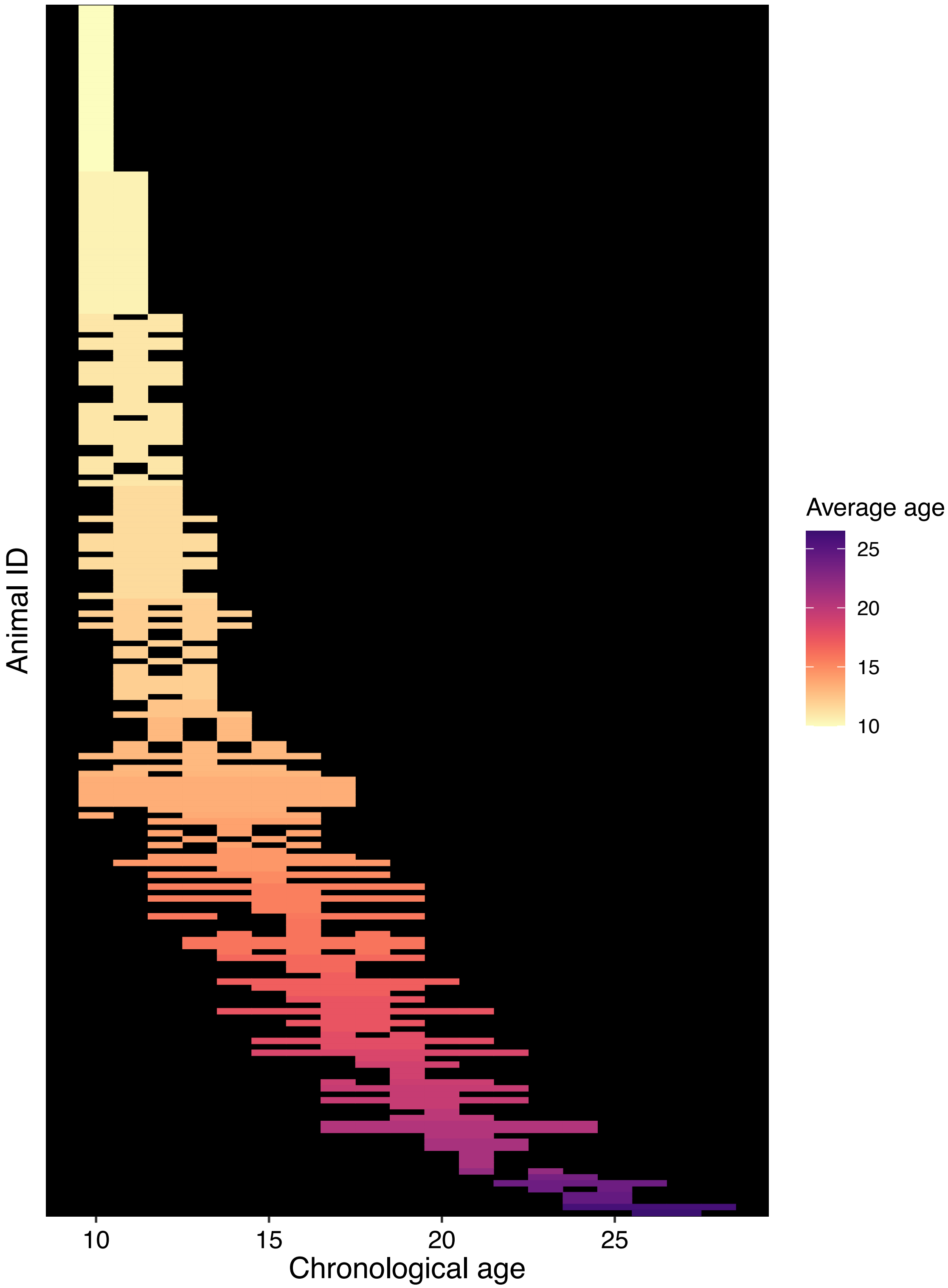


**Figure S1.** Heatmap showing age ranges over which each female macaque was sampled. Each row represents one individual.

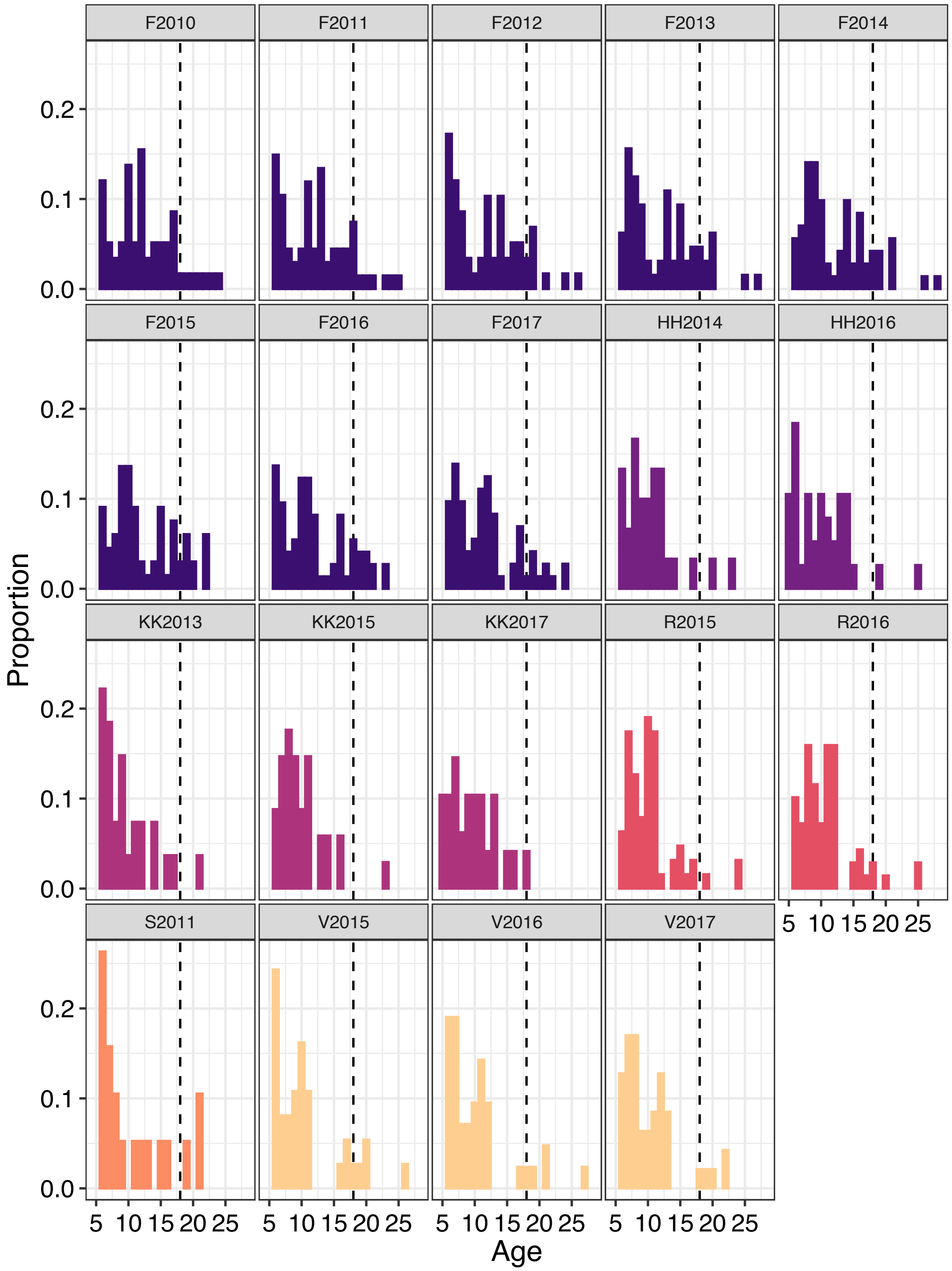


**Figure S2.** Distribution of ages in each of the 19 empirical macaque networks. The dotted black line at age 18 indicates the median age of death in this population and the cut off at which we considered individuals "old" for the sake of calculating the proportion of old individuals in the group.

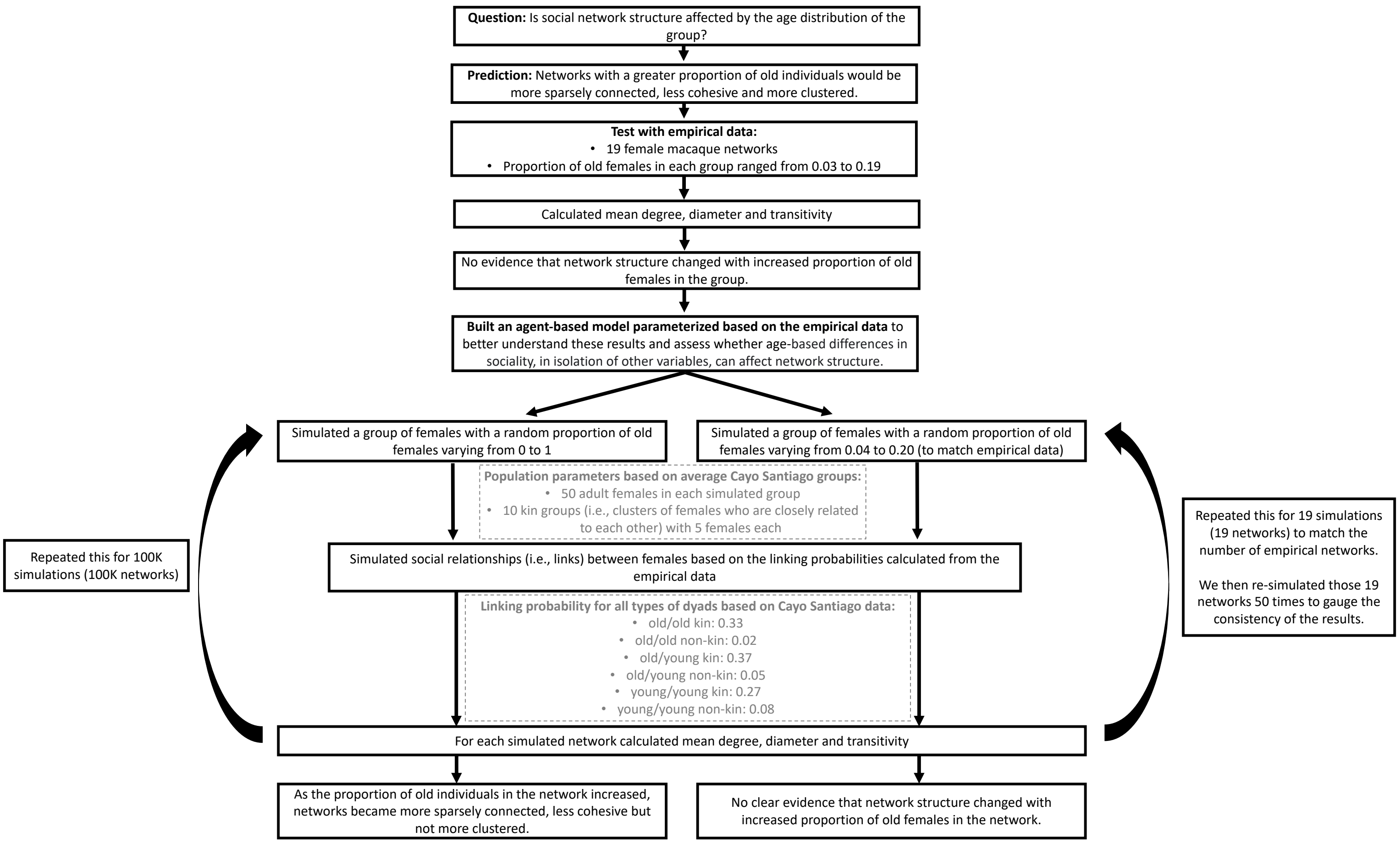


**Figure S3.** Schematic detailing the workflow of the agent-based model and the parameters used at various stages of the model-building process.

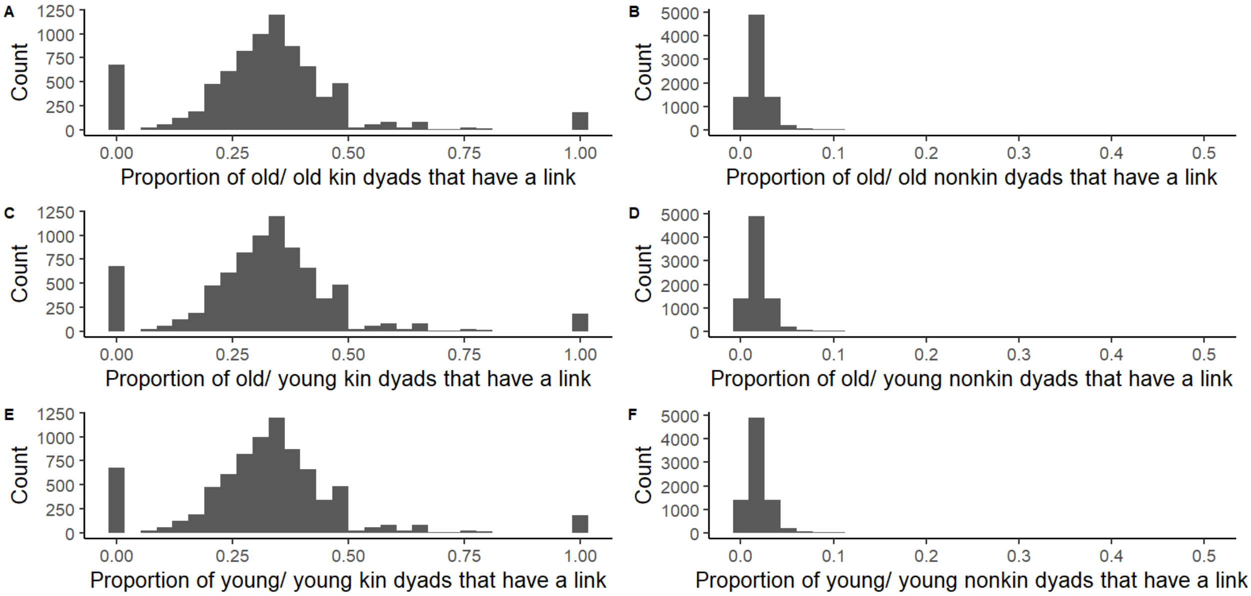


**Figure S4.** Results from 10000 simulated networks from the agent-based model showing the mean proportion of dyads from each age-kin class with a link. The mean proportion of dyads with a link was (A) 0.33 for old/old kin dyads, (B) 0.02 for old/old non-kin dyads, (C) 0.37 for old/young kin dyads, (D) 0.05 for old/young non-kin dyads, of (E) 0.27 for young/young kin dyads and (F) 0.08 for young/young non-kin dyads.

**Table S1.** Linking probabilities for the different types of dyads for all convergence steps. For each convergence step, each linking probability is equally distanced from the previous step and the next step. For each set of linking probabilities, we ran the model 10000 times.

| **Convergence step** | **Old/old kin** | **Old/old non-kin** | **Old/young kin** | **Old/young non-kin** | **Young/young kin** | **Young/young non-kin** |
|---|---|---|---|---|---|---|
| **1 (original probabilities)** | 0.330 | 0.020 | 0.370 | 0.050 | 0.270 | 0.080 |
| **2** | 0.325 | 0.025 | 0.358 | 0.050 | 0.275 | 0.075 |
| **3** | 0.320 | 0.030 | 0.347 | 0.050 | 0.280 | 0.070 |
| **4** | 0.315 | 0.035 | 0.335 | 0.050 | 0.285 | 0.065 |
| **5** | 0.310 | 0.040 | 0.323 | 0.050 | 0.290 | 0.060 |
| **6** | 0.305 | 0.045 | 0.312 | 0.050 | 0.295 | 0.055 |
| **7 (convergence point)** | 0.300 | 0.050 | 0.300 | 0.050 | 0.300 | 0.050 |

**Table S2.** Results from Bayesian mixed-effects model with a Gaussian error distribution looking at the **effect of age on grooming eigenvector centrality**. Response variable was logit transformed prior to analysis. Bolded terms indicate fixed effects where the 95% credible intervals did not overlap zero, providing evidence that those effects were significantly different from zero.

We fitted the model with the following weakly informative prior means and standard deviations (μ, σ): intercept (-2, 1), within-age (0, 0.5), average-age (0, 0.5), rankH (0, 0.5), rankM (0, 0.5).

| Effect | Group | Term | Estimate | Lower 95% CI | Upper 95% CI |
|---|---|---|---|---|---|
| **Fixed Effects** | | intercept | -1.37 | -3.22 | 0.58 |
| | | within-age | -0.04 | -0.30 | 0.24 |
| | | **average-age** | **-0.13** | **-0.22** | **-0.05** |
| | | rankM | 0.06 | -0.46 | 0.57 |
| | | **rankH** | **1.04** | **0.43** | **1.63** |
| **Random Effects** | Group | sd(intercept) | 2.26 | 1.08 | 4.44 |
| | Individual.ID | sd(intercept) | 0.64 | 0.05 | 1.19 |
| | | sd(within-age) | 0.17 | 0.01 | 0.44 |
| | | cor(intercept,within-age) | 0.36 | -0.86 | 0.98 |
| | Residual | sd(observation) | 3.09 | 2.87 | 3.31 |
| | Year | sd(intercept) | 1.02 | 0.40 | 2.10 |

**Table S3.** Results from Bayesian mixed-effects model with a zero-inflated Poisson error distribution looking at the **effect of age on grooming betweenness centrality**. Bolded terms indicate fixed effects where the 95% credible intervals did not overlap zero, providing evidence that those effects were significantly different from zero.

We fitted the model with the following weakly informative prior means and standard deviations (μ, σ): intercept (5, 1), within-age (0, 0.2), average-age (0, 0.2), rankH (0, 0.2), rankM (0, 0.2), within-age:rankH (0, 0.2), within-age:rankM (0, 0.2).

| Effect | Group | Term | Estimate | Lower 95% CI | Upper 95% CI |
|---|---|---|---|---|---|
| **Fixed Effects** | | intercept | 4.28 | 3.30 | 5.36 |
| | | within-age | 0.07 | -0.11 | 0.25 |
| | | average-age | -0.05 | -0.09 | 0.00 |
| | | **rankM** | **0.87** | **0.78** | **0.95** |
| | | **rankH** | **0.96** | **0.85** | **1.07** |
| | | **within-age:rankH** | **-0.35** | **-0.41** | **-0.29** |
| | | **within-age:rankM** | **-0.14** | **-0.18** | **-0.09** |
| **Random Effects** | Group | sd(intercept) | 0.84 | 0.29 | 2.05 |
| | Individual.ID | sd(intercept) | 1.03 | 0.91 | 1.18 |
| | | sd(within-age) | 1.04 | 0.90 | 1.20 |
| | | cor(intercept,within-age) | 0.11 | -0.11 | 0.32 |
| | Year | sd(intercept) | 0.24 | 0.12 | 0.49 |

**Table S4.** Results from Bayesian mixed-effects model with a Beta error distribution looking at the **effect of age on grooming closeness centrality**. Bolded terms indicate fixed effects where the 95% credible intervals did not overlap zero, providing evidence that those effects were significantly different from zero.

We fitted the model with the following weakly informative prior means and standard deviations (μ, σ): intercept (-3, 2), within-age (0, 1), average-age (0, 1), rankH (0, 1), rankM (0, 1).

| Effect | Group | Term | Estimate | Lower 95% CI | Upper 95% CI |
|---|---|---|---|---|---|
| **Fixed Effects** | | intercept | -3.93 | -5.46 | -2.29 |
| | | **within-age** | **-0.18** | **-0.35** | **-0.02** |
| | | average-age | 0.00 | -0.03 | 0.02 |
| | | **rankM** | **0.17** | **0.04** | **0.30** |
| | | rankH | 0.15 | -0.02 | 0.33 |
| **Random Effects** | Group | sd(intercept) | 1.27 | 0.63 | 2.61 |
| | Individual.ID | sd(intercept) | 0.57 | 0.49 | 0.66 |
| | | sd(within-age) | 0.87 | 0.70 | 1.05 |
| | | cor(intercept,within-age) | 0.59 | -0.01 | 0.82 |
| | Year | sd(intercept) | 1.76 | 1.02 | 3.12 |

**Table S5.** Results from Bayesian mixed-effects model with a zero-one-inflated Beta error distribution looking at the **effect of age on grooming clustering coefficient**. Bolded terms indicate fixed effects where the 95% credible intervals did not overlap zero, providing evidence that those effects were significantly different from zero.

We fitted the model with the following weakly informative prior means and standard deviations (μ, σ): intercept (-2, 1), within-age (0, 1), average-age (0, 1), rankH (0, 1), rankM (0, 1).

| Effect | Group | Term | Estimate | Lower 95% CI | Upper 95% CI |
|---|---|---|---|---|---|
| **Fixed Effects** | | intercept | -1.53 | -2.02 | -1.06 |
| | | within-age | 0.04 | -0.06 | 0.14 |
| | | average-age | 0.03 | 0.00 | 0.06 |
| | | rankM | 0.06 | -0.16 | 0.29 |
| | | rankH | 0.15 | -0.08 | 0.39 |
| **Random Effects** | Group | sd(intercept) | 0.11 | 0.00 | 0.40 |
| | Individual.ID | sd(intercept) | 0.13 | 0.01 | 0.31 |
| | | sd(within-age) | 0.21 | 0.11 | 0.33 |
| | | cor(intercept,within-age) | 0.27 | -0.81 | 0.96 |
| | Year | sd(intercept) | 0.18 | 0.01 | 0.51 |

**Table S6.** Results from Bayesian mixed-effects model with a Gaussian error distribution looking at how **mean degree of grooming networks** changes with the proportion of old individuals in the group. Bolded terms indicate fixed effects where the 95% credible intervals did not overlap zero, providing evidence that those effects were significantly different from zero.

We fitted the model with the following weakly informative prior means and standard deviations (μ, σ): intercept (4, 1), proportion old (0, 5), average relatedness (0, 5).

| Effect | Group | Term | Estimate | Lower 95% CI | Upper 95% CI |
|---|---|---|---|---|---|
| **Fixed Effects** | | intercept | 4.18 | 2.95 | 5.44 |
| | | proportion old | -2.76 | -10.24 | 5.25 |
| | | average relatedness | -0.54 | -10.10 | 8.94 |
| **Random Effects** | Year | sd(intercept) | 0.74 | 0.04 | 1.98 |
| | Residual | sd(observation) | 1.41 | 0.96 | 2.07 |

**Table S7.** Results from Bayesian mixed-effects model with a Gaussian error distribution looking at how **diameter of grooming networks** changes with the proportion of old individuals in the group. Response variable was log transformed prior to analysis. Bolded terms indicate fixed effects where the 95% credible intervals did not overlap zero, providing evidence that those effects were significantly different from zero.

We fitted the model with the following weakly informative prior means and standard deviations (μ, σ): intercept (5.5, 0.5), proportion old (0, 2.5), average relatedness (0, 2.5), density (0, 2.5).

| Effect | Group | Term | Estimate | Lower 95% CI | Upper 95% CI |
|---|---|---|---|---|---|
| **Fixed Effects** | | intercept | 5.68 | 4.91 | 6.42 |
| | | proportion old | -0.12 | -3.93 | 3.67 |
| | | average relatedness | 0.05 | -4.76 | 4.79 |
| | | density | -3.65 | -7.26 | 0.23 |
| **Random Effects** | Year | sd(intercept) | 0.44 | 0.02 | 1.16 |
| | Residual | sd(observation) | 0.66 | 0.43 | 1.00 |

**Table S8.** Results from Bayesian mixed-effects model with a Gaussian error distribution looking at how **transitivity of grooming networks** changes with the proportion of old individuals in the group. Bolded terms indicate fixed effects where the 95% credible intervals did not overlap zero, providing evidence that those effects were significantly different from zero.

We fitted the model with the following weakly informative prior means and standard deviations (μ, σ): intercept (0.15, 0.1), proportion old (0, 1), average relatedness (0, 1), density (0, 1).

| Effect | Group | Term | Estimate | Lower 95% CI | Upper 95% CI |
|---|---|---|---|---|---|
| **Fixed Effects** | | intercept | 0.09 | -0.01 | 0.19 |
| | | proportion old | 0.20 | -0.29 | 0.70 |
| | | average relatedness | -0.66 | -2.35 | 1.06 |
| | | **density** | **0.81** | **0.38** | **1.22** |
| **Random Effects** | Year | sd(intercept) | 0.04 | 0.00 | 0.09 |
| | Residual | sd(observation) | 0.05 | 0.03 | 0.08 |

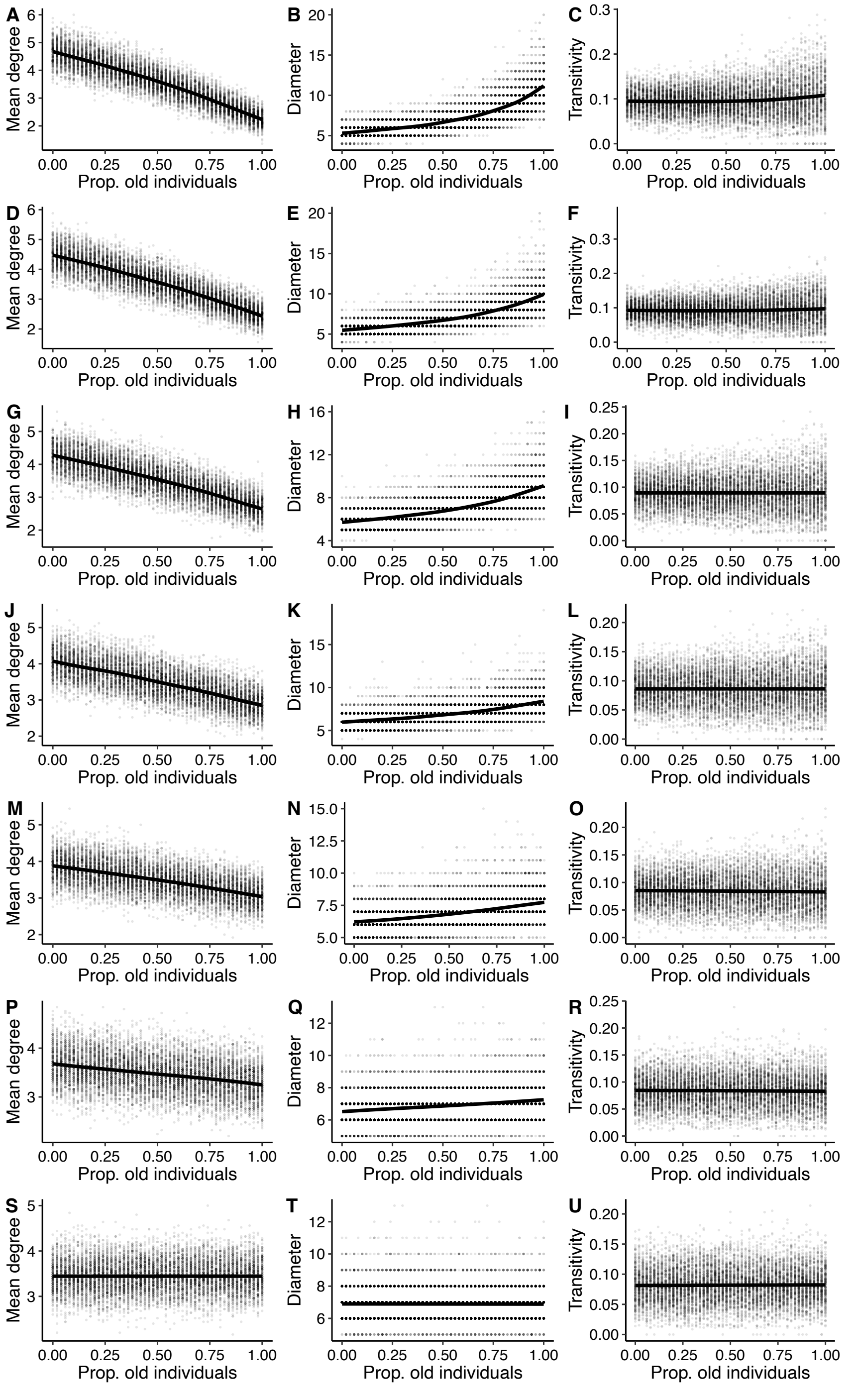


**Figure S5.** Effects of the proportion of old individuals on network structure under reduction of the difference in linking probabilities between the two age categories (old and young). The difference was reduced in 7 convergence steps and linking probabilities for each step are given in Table S1 above. The convergence steps shown in the figure are as follows: (**A-C**) convergence step 1 (original probabilities); (**D-F**) convergence step 2; (**G-I**) convergence step 3; (**J-L**) convergence step 4; (**M-O**) convergence step 5; (**P-R**) convergence step 6; (**S-U**) convergence step 7 (no difference between age categories).

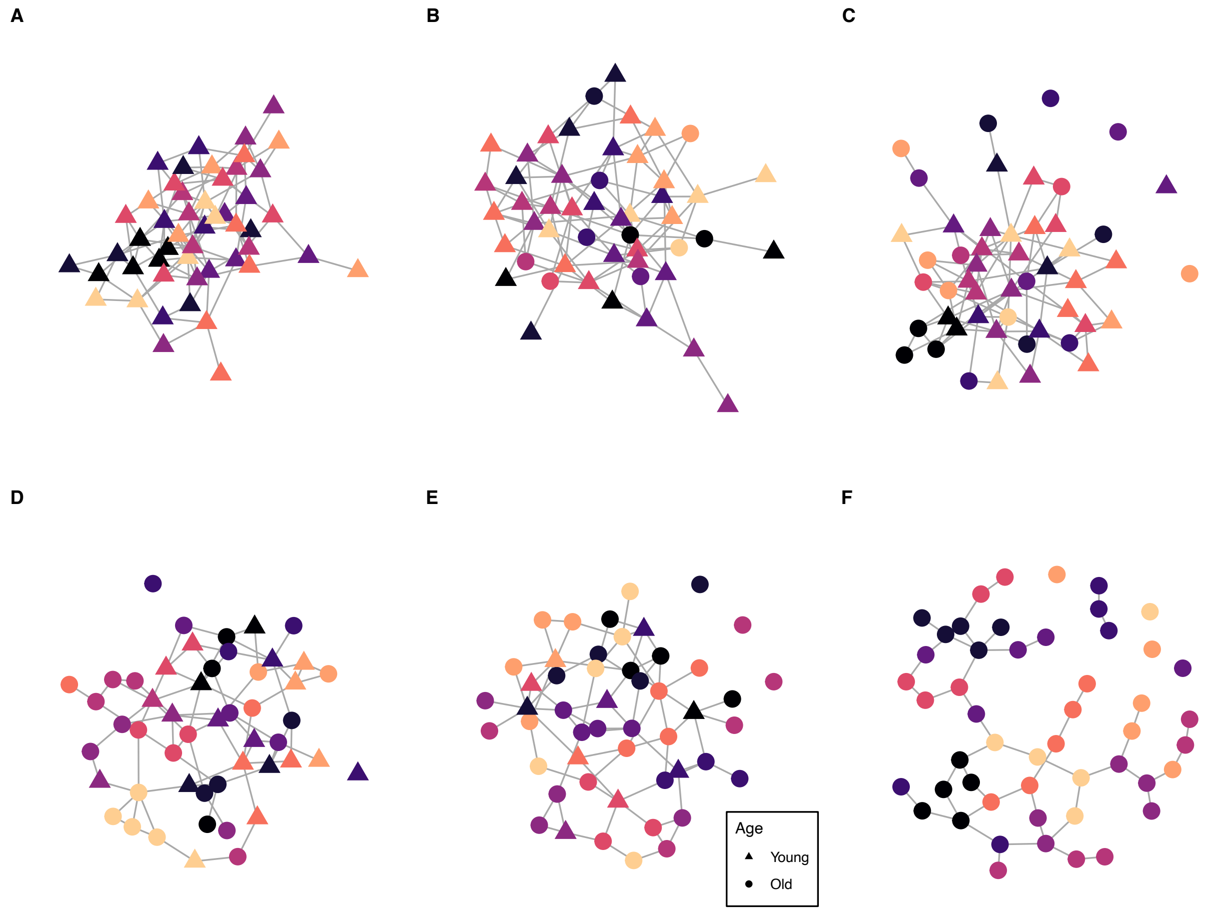


**Figure S6**. Example networks from the agent-based model illustrating differences in network structure with different proportions of old individuals in the network. Proportions of old individuals are as follows: (**A**) 0, (**B**) 0. 2, (**C**) 0.4, (**D**), 0.6, (E) 0.8, (**F**) 1.
